# TrmB family transcription factor as a thiol-based regulator of oxidative stress response

**DOI:** 10.1101/2022.03.05.483106

**Authors:** Paula Mondragon, Sungmin Hwang, Lakshmi Kasirajan, Rebecca Oyetoro, Angelina Nasthas, Emily Winters, Ricardo L. Couto-Rodriguez, Amy Schmid, Julie A. Maupin-Furlow

## Abstract

Oxidative stress causes cellular damage including DNA mutations, protein dysfunction and loss of membrane integrity. Here we discovered TrmB (transcription regulator of *mal* operon) family proteins (Pfam PF01978) composed of a single winged-helix DNA binding domain (InterPro IPR002831) can function as thiol-based transcriptional regulators of oxidative stress responses. Using the archaeon *Haloferax volcanii* as a model system, we demonstrate that the TrmB-like OxsR is important for recovery of cells from hypochlorite stress. OxsR is shown to bind specific regions of genomic DNA, particularly during hypochlorite stress. OxsR-bound intergenic regions were found proximal to oxidative stress operons including genes associated with thiol relay and low molecular weight thiol biosynthesis. Further analysis of a subset of these sites, revealed OxsR to function during hypochlorite stress as a transcriptional activator and repressor. OxsR was shown to require a conserved cysteine (C24) for function and to use a CG-rich motif upstream of conserved BRE/TATA box promoter elements for transcriptional activation. Protein modeling suggested the C24 is located at a homodimer interface formed by antiparallel α helices, and that oxidation of this cysteine would result in the formation of an intersubunit disulfide bond. This covalent linkage may promote stabilization of an OxsR homodimer with the enhanced DNA binding properties observed in the presence of hypochlorite stress. The phylogenetic distribution TrmB family proteins, like OxsR, that have a single winged-helix DNA binding domain and conserved cysteine residue suggests this type of redox signaling mechanism is widespread in Archaea.

**Importance:** TrmB-like proteins, while not yet correlated with redox stress, are found in bacteria and widespread in archaea. Here we expand annotation of a large group of TrmB-like single winged-helix DNA binding domain proteins from diverse archaea to function as thiol-based transcriptional regulators of oxidative stress response. Using *Haloferax volcanii* as a model, we reveal the TrmB-like OxsR functions during hypochlorite stress as a transcriptional activator and repressor of an extensive gene co-expression network associated with thiol relay and other related activities. A conserved cysteine residue of OxxR serves as the thiol-based sensor for this function and likely forms an intersubunit disulfide bond during hypochlorite stress that stabilizes a homodimeric configuration with enhanced DNA binding properties. A CG-rich DNA motif in the promoter region of a subset of sites identified to be OxsR-bound is required for regulation; however, not all sites have this motif suggesting added complexity to the regulatory network.

## Introduction

Oxidative stress can be exceedingly damaging to cells. Once the levels of reactive species overwhelm the antioxidant capacity of cells, lipid peroxidation, protein denaturation, DNA hydroxylation, and other damaging effects occur that impair cellular viability (1). To survive these challenges, cells must sense and respond to oxidant challenge, shifts in redox balance, and the damage encountered due to oxidative stress. Transcription factors (TFs) that sense and respond to reactive species are found to mediate global changes in gene expression to remedy the damage (2, 3). These TFs often have metalloclusters, cofactors, or residues (cysteine, histidine or methionine) that are sensitive to oxidant and serve as “redox switches” (4, 5). These switches can turn TF function on or off in coordinating gene co-expression networks associated with electron flow, antioxidant systems and other pathways (6–8).

Archaea have an array of metabolic strategies and physiological adaptations that enable them to sense and respond to extreme environments. TFs are central to sensing and responding to environmental cues and in relaying this signal to coordinate gene-expression networks. Thus, TFs are anticipated to be important in facilitating the ability of archaea to respond and thrive in extreme environments including exposure to reactive species (9, 10). TFs with redox switches are diverse and widespread in bacteria and likely to function in archaea to sense oxidant rich conditions. Archaea, while distinct from bacteria in their use a eukaryotic-like basal transcription machinery, have TFs that are often related to bacteria (9, 10). This similarity in TFs is due to their evolution which includes shared ancestral proteins and inter-domain horizontal gene transfer events between archaea and bacteria. Thus, it is surprising that the bacterial TFs, such as OhrR (11–13), SarA/MgrA (14), PerR (15), HypR (16), YodB (17), QsrR (18), MosR (19), SarZ (20), OxyR (21–23), SoxR (24, 25), and FNR (26), that use redox switches to mediate global alterations of transcriptional networks are not readily identified in archaea.

The two TFs that have been identified in archaea to use redox switches are from the ArsR family, *i.e*., MsvR (MTH_1349) and SurR (PF0095). Oxidation of key cysteine residues of these TFs negatively affects their DNA binding activity. SurR responds to elemental sulfur (S^0^), an electron acceptor in the *Thermococcales*, leading to the deactivation of genes associated with H_2_ production and the derepression of genes needed for S^0^ metabolism (27–30). MsvR regulates the transcription of its own promoter and an adjacent operon implicated in the oxidative stress response of methanogens (31–33). MsvR appears exclusive to methanogens, while SurR clusters to the Archaeal Clusters of Orthologous Gene (arCOG) group arCOG01684 (34), suggesting its general function is more widespread.

TrmB (transcription regulator of *mal* operon) family TFs, though not yet correlated with redox stress, are widespread in archaea and found in bacteria (9, 35). TrmB family proteins appear to have undergone an evolutionary expansion in archaea after divergence from bacteria, with homologs accounting for 12% of the total number of TFs in archaea compared to 0.5% in bacteria (9). TrmB family TFs have an N-terminal DNA binding domain that is sometimes fused to a C-terminal ligand sensing domain. The ligands that bind to the C-terminal domain can be sugars or other molecular factors. TrmB family TFs with these two domains (the N-terminal DNA binding domain and C-terminal ligand sensing domain) typically function as global transcriptional activators and/or repressors of sugar transport and metabolism including glycolysis, gluconeogenesis, the TCA cycle, amino acid metabolism, methanogenesis, and autotrophic pathways (36–46). TrmB homologs with the single DNA binding domain, while less characterized, appear to function differently than their two-domain counterparts. This variation is exemplified by TrmBL2 (TrmB-like protein 2), an abundant DNA binding protein of the *Thermococcales* that clusters to arCOG02037. TrmBL2 functions as a global transcriptional repressor and a chromatin binding protein that can rearrange genomic DNA from a conventional histone-bound ‘beads-on-a-string’ architecture to a thick fibrous structure (34). When compared to other TrmB family proteins, TrmBL2 has an expanded function as it can bind single-stranded and double stranded DNA (47).

Identification of TFs and other global regulatory systems used by archaea to sense and respond to reactive species is imperative to provide new insights into hypertolerance mechanisms. Haloarchaea provide a useful resource for this discovery as they inhabit some of the saltiest places on Earth, including hypersaline lakes, marine salterns and brine inclusions, which are high in reactive species (48, 49). Compared to most organisms, haloarchaea display an order of magnitude higher tolerance to oxidant rich conditions, which is likely associated with their adaptation to these hypersaline ecosystems (50, 51). Haloarchaea employ an unusual ‘high-salt-in’ strategy in which molar concentrations of potassium and chloride are accumulated intracellularly to maintain osmoprotection and cellular homeostasis (52–55). These high concentrations of chloride promote the generation of reactive chloride species (RCS), reactive oxygen species (ROS), and other stressful agents (56–58). High levels of reactive species are also generated in hypersaline habitats through common cycles of desiccation-rehydration and intense UV radiation (56, 57, 59). Surprisingly, haloarchaea thrive under these conditions and are often the last remaining communities when the salt concentrations reach saturation (60).

Regulators of oxidative stress responses are identified in haloarchaea; however, the mechanisms of how these regulators sense oxidant remain to be determined. Included among these factors are RosR (VNG0258H) and SHOxi. RosR, a PadR-type TF with a winged helix-turn-helix (wHTH) domain, is required for gene expression dynamics during extreme oxidative stress in *Halobacterium* sp. NRC-1 (61–63). RosR homologs cluster to the arCOG00006 group and are found widespread among haloarchaea suggesting a common mechanism, yet the residues that may directly sense oxidant are not readily identifiable. SHOxi is a small non-coding RNA in *Haloferax volcanii* that impacts redox balance by destabilizing malic enzyme mRNA and, thus, decreasing the ratio of NAD^+^/NADH (64). SHOxi is upregulated during oxidative stress, suggesting other factors serve upstream of this response to sense the redox status of cells.

Here we discover that TrmB family proteins can function as TFs that sense redox status and facilitate the recovery of cells from hypochlorite stress. Using *Haloferax volcanii* as a model system, we targeted the TrmB family protein HVO_2970 of arCOG02242 for study, as it undergoes a several-fold increase in protein abundance after exposure of cells to hypochlorite stress as determined by stable isotope labelling of amino acids in cell culture (SILAC)-based proteomic analysis (65). Here we reveal HVO_2970 is a thiol-based TF required for recovery of cells from hypochlorite stress and, thus, it is named OxsR, for <u>ox</u>idative <u>s</u>tress responsive regulator. By coupling chromatin immunoprecipitation sequencing (ChIP-seq) and quantitative real-time polymerase chain reaction (qRT-PCR) analyses, we demonstrate that OxsR regulates the expression of genes associated with thiol relay, low molecular weight thiol synthesis, and other related functions during hypochlorite stress. OxsR functions as a transcriptional activator and repressor, with these distinctions found to correlate with the presence and positioning of a CG-rich motif relative to the BRE/TATA box promoter consensus sequence. By site-directed mutagenesis, a cysteine residue (C24) oriented at a predicted homodimer interface was demonstrated to be important for OxsR function. This cysteine residue was conserved among TrmB-like single DNA binding domain proteins from diverse archaeal phyla suggesting this type of redox signaling mechanism is widespread and represents a new type of thiol-based TF.

## Results

### OxsR phylogenetic distribution

OxsR is member of the TrmB family (Pfam PF01978). Like most archaea, *H. volcanii* encodes multiple TrmB family proteins with all thirteen harboring an N-terminal DNA binding domain (IPR002831) classified to the winged helix-turn-helix (wHTH) superfamily (IPR036388) (**Figure 1A**). However, only five of these proteins were fused to a C-terminal ligand sensing domain (IPR021586). The remaining eight proteins had a single DNA binding domain including the smallest member of this group: OxsR (123 aa, 14 kDa). Such single-domain TrmB family proteins are widespread among archaeal phyla yet poorly understood in function, while two-domain TrmB homologs with a C-terminal ligand binding domain are more commonly studied but not as widely distributed (**Figure 1B**). Within the TrmB family, OxsR is classified to the archaeal cluster of orthologous genes arCOG02242 (66) and by BlastP comparison is related to homologs of the *Euryarchaeota*, *Crenarchaeota* and *Asgard* archaea (**Figure 1B**). OxsR does not share close relationship to bacterial members of the TrmB family, which are less common than those of archaea (9). Overall, OxsR is a TrmB family protein that has a single wHTH domain and appears conserved in diverse archaeal phyla.

**Figure 1.**
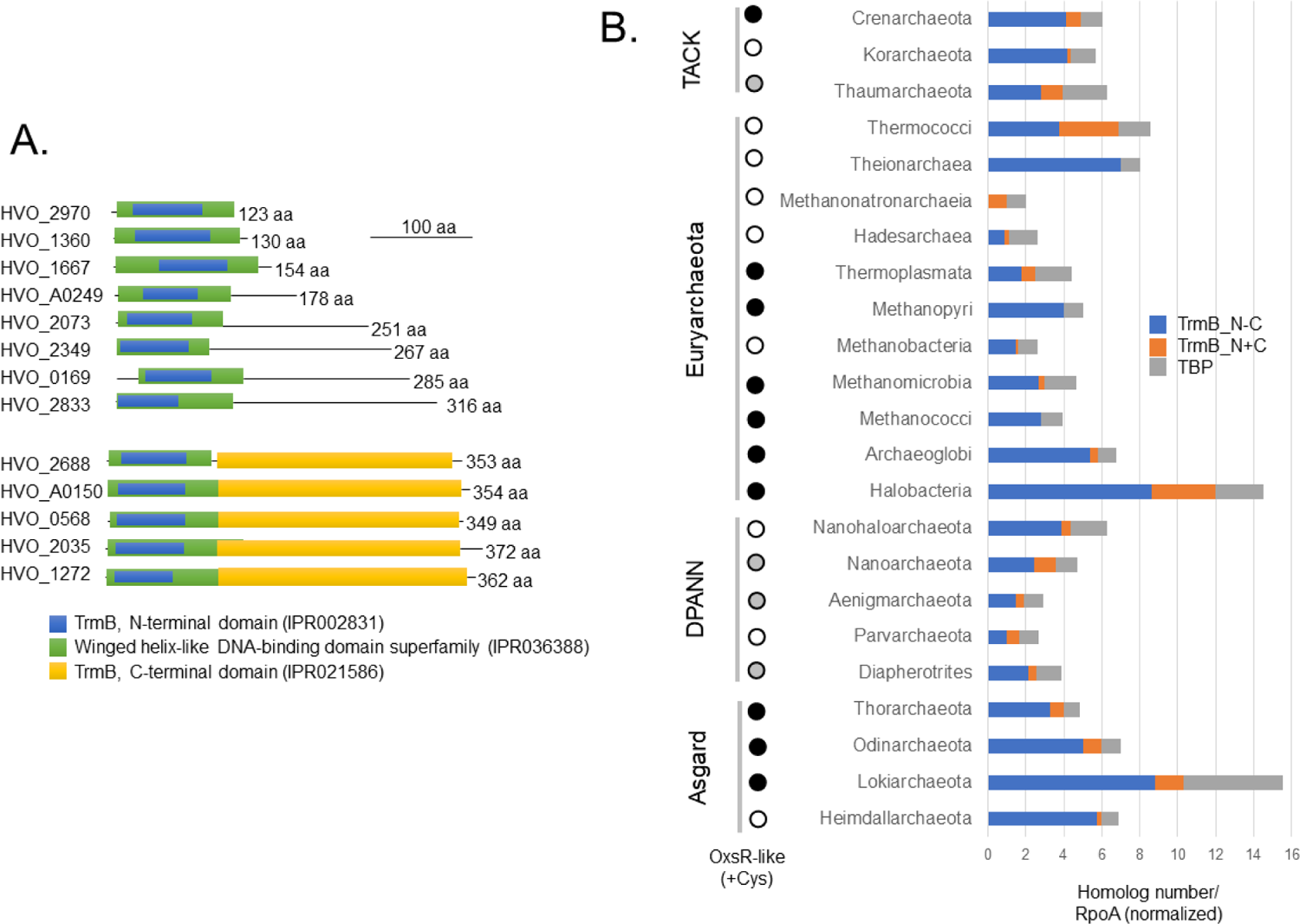
*Haloferax volcanii* OxsR (HVO_2970) is a member of the TrmB family and is conserved across multiple phyla of *Archaea*. A) Two major types of TrmB family members observed. *H. volcanii* TrmB family proteins with: i) an N-terminal winged helix-turn-helix (wHTH) DNA binding domain (green and blue) and a C-terminal ligand (sugar) binding domain (yellow) and ii) only the wHTH domain. OxsR is of the latter type. B) Distribution of OxsR and TrmB family homologs among archaea. TrmB family proteins with only an N-terminal DNA binding domain (IPR002831) (blue bar), TrmB proteins with an N-terminal DNA binding domain and C-terminal ligand sensing domain (IPR021586) (orange bar), and TATA binding proteins (TBPs, IPR000814) (grey bar) are compared as ratios of UniProt hits per RNA polymerase alpha subunit (RpoA, IPR000722) to account for differences in genome sequence availability among the phyla. Among the archaea, TrmB homologs with a C-terminal ligand sensing domain were not detected in *Methanopyri*, *Methanococci* and *Theionarchaea*. Archaea with cysteine-containing OxsR homologs are indicated by black and grey circles (with grey indicating that many archaea within this group do not contain OxsR homologs).

### OxsR is important for recovery of *H. volcanii* from hypochlorite stress

To examine the biological role of OxsR, the following strains were constructed. The *oxsR* gene was deleted from the *H. volcanii* genome (H26 parent) by markerless deletion (**Figure S1, Table S1**). The resulting *ΔoxsR* mutant was subsequently transformed with plasmids carrying the *oxsR* gene and the empty vector control. In addition, a strain for OxsR pull-down assays was generated by integration of the hemagglutinin (HA) coding sequence at the 3’ end of *oxsR* on the genome.

The *oxsR*-HA integrant strain was also modified to include a C24A mutation discussed in a later section. The strains constructed in this manner were subsequently examined for recovery from hypochlorite stress. All strains were found to be of comparable growth rate (1.7-1.9 h generation time) when cultured under standard conditions in glycerol minimal medium (GMM) (**Figure 2A**).

**Figure 2.**
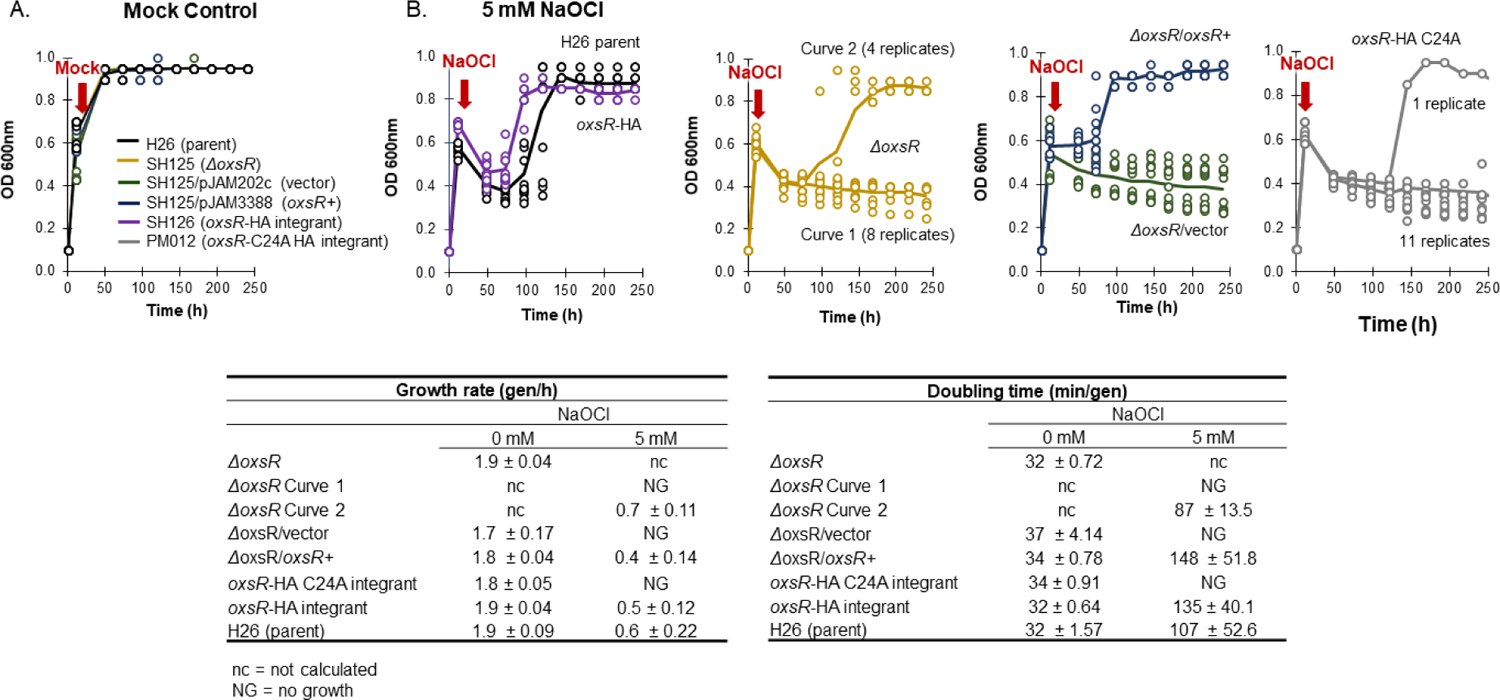
OxsR facilitates recovery of *H. volcanii* from hypochlorite stress. Strains were grown glycerol minimal medium (GMM) to log-phase (11 hr, OD600 nm, 0.5-0.6) and then treated with a mock control (A) or 5 mM NaOCl (B) as indicated. Lower panels, Growth rates (left) and doubling times (right) are given for each strain under optimal conditions (0 mM) or hypochlorite stress (5 mM NaOCl). The markers represent the individual replicates (n = 12 total; 6 replicates x 2 experiments). The colored curves represent the average growth for the 12 replicates with exception of the NaOCl treated *ΔoxsR* mutant and *oxsR*-HA C24A integrant strains which are represented by two curves. Cell growth was monitored at OD600 nm by direct measurement in the 13 x 100 mm culture tubes using a Spectronic 20+ spectrophotometer (ThermoSpectronic, Filter: 600-950 nm). The measurement plateau at OD600 of 0.95 is the maximum OD600 the spectrophotometer could read. See methods for details.

By contrast, differences in strain recovery were observed after hypochlorite treatment (**Figure 2B**). The parent and HA-tagged strains were found to fully recover from hypochlorite stress after an 88 ± 8 h lag (**Figure 2B**). By contrast, the *ΔoxsR* mutant displayed no growth (curve 1, 8 replicates) or recovered after an extended lag (117 ± 23 h) (curve 2, 4 replicates) under these same conditions (**Figure 2B**). All replicates of the mutant retained the markerless deletion of the *oxsR* gene, consistent with whole genome resequencing analysis which revealed *oxsR* to be deleted from all genomic copies (**Figure S2A-C, Table S1**). One distinction was identified among the *ΔoxsR* mutant replicates: those that that did not recover from hypochlorite stress initially (curve 1) were no longer viable, whereas the replicates that displayed an extended lag (curve 2) were viable, but no longer able to recover after exposure to a second dose of hypochlorite and were not viable once this stress agent was removed (**Figure S2D**). When performing complementation assays, the *ΔoxsR* mutant was found to be uniformly hypersensitive to hypochlorite in the presence of the empty vector, suggesting that the plasmid posed an extra burden to the cells under these conditions (**Figure 2B**). Expression of the *oxsR* gene from this multicopy plasmid restored the *ΔoxsR* mutant to recover from the hypochlorite stress at parental levels (**Figure 2B**). This finding suggests that the second site point mutation observed in the *ΔoxsR* genome sequence (non-coding intergenic G>A mutation between *hvo_RS01570* and *hvo_RS01575,* **Table S1**) was not the source of the observed hypochlorite recovery defect of *ΔoxsR*. Based on these results, OxsR is important for the recovery of *H. volcanii* from hypochlorite stress. Furthermore, the minimal effect of the C-terminal HA-tag on OxsR function during hypochlorite stress (**Figure 2B**) indicated this construct could be used for downstream immunoprecipitation analysis.

### OxsR binds specific regions on the *H. volcanii* genome

OxsR was investigated for its ability to bind specific and/or nonspecific regions of *H. volcanii* genomic DNA by combining chromatin immunoprecipitation with massively parallel DNA sequencing (ChIP-seq). The parent (H26) and *oxsR-HA* tagged strains were grown to log-phase in GMM and then treated with hypochlorite or a mock control prior to the ChIP-seq analysis (see Methods for details). By this approach, OxsR-bound sites were investigated over the chromosome and endogenous plasmids. A total of 29 and 130 sites were found to be putative OxsR interacting regions in the absence and presence of oxidative stress, respectively (**Figure S3, Table S2**). Although many peaks (63%) were located within the coding sequences of genes, OxsR was also found to bind distinct intergenic regions of the genome. Most of the OxsR-bound intergenic sites (49 of 59 total) were detected only in the hypochlorite-treated cells with six operons having two 5’ binding sites. Of the remaining intergenic sites, seven were identified irrespective of the treatment and three were detected only in the absence of hypochlorite (**Table S2**). When analyzing the genes adjacent to the OxsR-bound regions by arCOG gene functional analysis (67), nearly half (44%, hypergeometric test *p*-value of enrichment relative to genomic background < 1.1 x 10^-2^) of the encoded proteins clustered to the arCOG [S] group of unknown function, thus, providing limited insight (**Figure 3**). However, 12% of the proteins clustered to the arCOG [O] group associated with post-translational modification, protein turnover, and chaperone functions, including gene homologs associated with thiol relay (*e.g*., thioredoxin, thioredoxin-like and disulfide oxidoreductase activity, **Table S2**); and 3% to [I] (lipid transport and metabolism, *p* < 6.1 x 10^-5^). Additional functional analysis of the ChIP-seq associated genes by STRING (11.5)(68) corroborated arCOG findings (false discovery rates < 0.0374; **Figure S4**). Further inspection of the sites strongly enriched for OxsR binding in intergenic regions (“ChIP-seq high peaks” with height > 1000-fold enrichment relative to the input control, 27 of 59 total, **Table S2**) revealed that at least seven of the linked genes were associated with the synthesis of low molecular weight thiols and thiol relay systems (**Table 1**). HVO_1043, a member of the DUF1684 family, was included in this thiol relay group, as 3D-modeling and multiple amino acid sequence alignment revealed a conserved Cx_7_C motif with the thiol groups of these cysteines in a proximity typical of function in thiol relay (**Figure S5**). Overall, these results reveal OxsR is a TF homolog that binds specific regions of genomic DNA during redox stress. Of the intergenic regions bound by OxsR, an enrichment in the promoter regions of genes associated with protein quality control, thiol relay and other related functions predicted to be important in the recovery from hypochlorite stress was observed.

**Figure 3.**
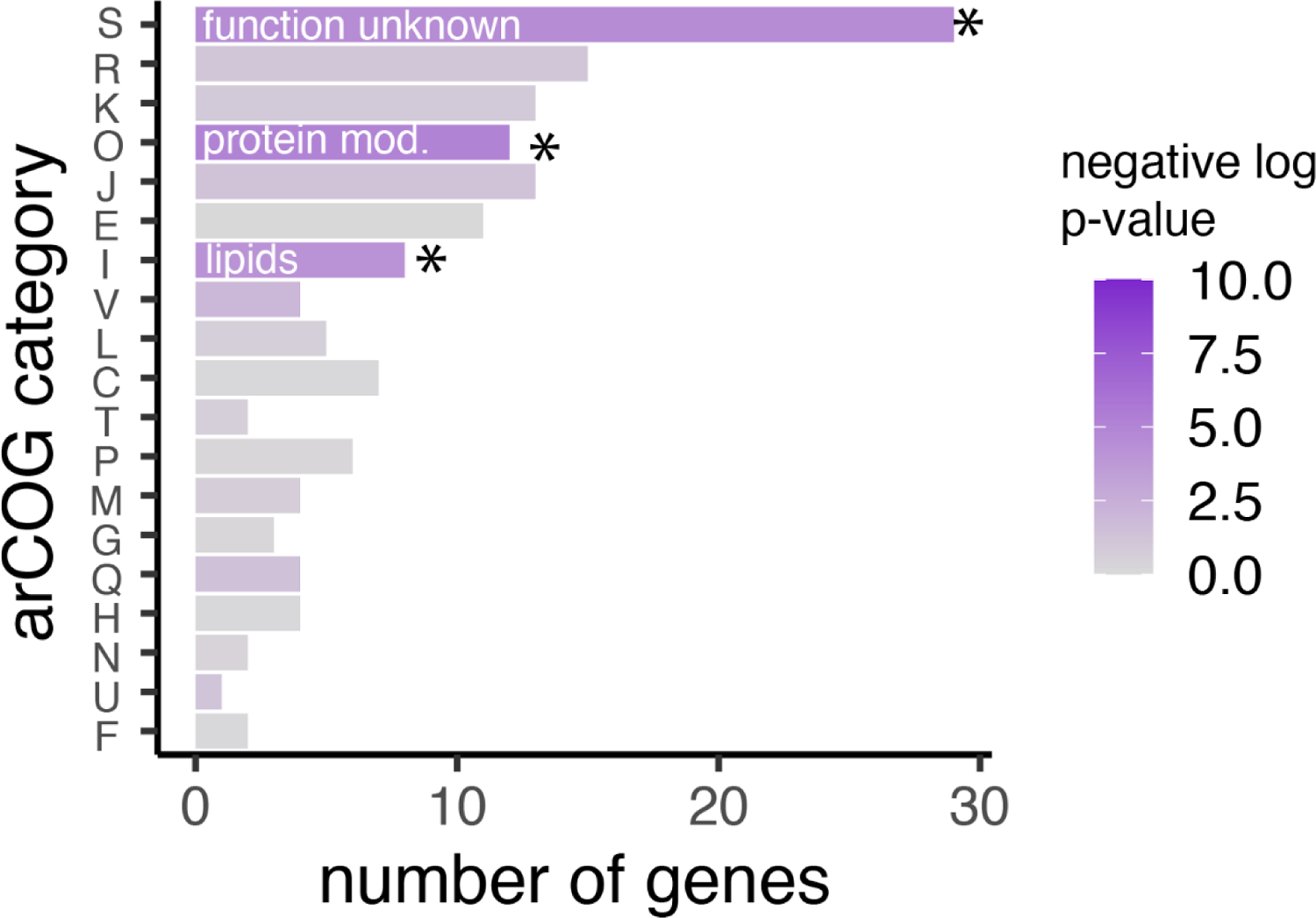
General functional classification of genes predicted to be bound by OxsR according to ChIP-seq data. Gene homologs flanking the intergenic regions of DNA detected to be bound by OxsR through ChIP-seq analysis were clustered by arCOG according to function. Bar lengths represent the number of genes detected in each functional category, and the shading of the bar represents the significance of enrichment in each category (see legend).

**Table 1.** Gene loci with 5’ regions detected as high peaks by OxsR ChIP seq analysis^1^.

| Gene locus | Protein names | NaOCl (-) | NaOCl (+) | High peak |
| --- | --- | --- | --- | --- |
| HVO_1043* | DUF1684 family protein | V | V | V |
| HVO_A0618* | Coenzyme A disulfidereductase homolog | V | V | V |
| HVO_1875 | Galactoside O-acetyltransferase (EC 2.3.1.-) homolog | V | V | V |
| HVO_0198 | UPF0213 family protein | V | V | V |
| HVO_1342* | Thioredoxin | V | V | V |
| HVO_0811* | L-Aspartate decarboxylase (EC 4.1.1.11) homolog |  | V | V |
| HVO_0337 | Glutaredoxin |  | V | V |
| HVO_0040* | Transmembrane protein |  | V | V |
| HVO_0004 | NamA family oxidoreductase |  | V | V |
| HVO_1031 | Thioredoxin-disulfide reductase (EC 1.8.1.9) |  | V | V |
| HVO_1166 | Chloroplast RNA splicing and ribosome maturation (CRM) domain protein |  | V | V |
| HVO_1684 | Threonine-tRNA ligase (EC 6.1.1.3) |  | V | V |
| HVO_0497 | Cold shock protein |  | V | V |
| HVO_0251 | Glutaredoxin |  | V | V |
| HVO_1244 | Thioredoxin |  | V | V |
| HVO_1668* | γ-Glutamylcysteine synthetase (EC 6.3.2.2) | | V | V |
| HVO_1522 | Hexuronic acid methyltransferase AgIP (EC 2.1.1.-) |  | V | V |
| HVO_1438* | YneT family protein |  | V | V |
| HVO_1457 | Glycoside hydrolase domain protein |  | V | V |
| HVO_2157 | Uncharacterized protein |  | V | V |
| HVO_2859 | DUF63 family protein |  | V | V |
| HVO_1124 | DUF357 family protein |  |  | V |
| HVO_1017 | Ketosamine kinase domain protein |  |  | V |

<sup>1</sup>Blue highlight, putative function in low molecular weight thiol (LMWT) biosynthesis or thiol relay; \*, CG-rich DNA motif CGGnCGnGCG identified 5' of these gene homologs. V, present in the condition.

### OxsR functions as an apparent transcription factor

Five genes were selected from the ChIP-seq high peak group to determine whether OxsR has an impact on their transcript abundance. In addition, *oxsR* (*hvo_2970*) was included to examine how its transcript levels may correlate with the 3-fold increase in OxsR protein abundance previously identified during hypochlorite stress (69). Transcript levels were monitored by real-time quantitative reverse transcription polymerase chain reaction (RT-qPCR) in the parent and *ΔoxsR* mutant after treatment with hypochlorite or a mock control (t = 0) (**Figure 4**). Four of the five genes examined from the ChIP-seq dataset were found to be upregulated in the parent after exposure to hypochlorite including the transcripts of: i) *hvo_0811* and *hvo_1043*, which displayed a rapid increase; and ii) *hvo_0040* and *hvo_0337,* which were more delayed in their upregulation. The exception was *hvo_0039*, whose expression remained relatively constant in the parent strain regardless of time or treatment. Hypochlorite stress was also found to stimulate a transient increase in the *oxsR* transcript levels which may in part account for the 3-fold increase in OxsR protein abundance. When comparing the transcript profiles, the *ΔoxsR* mutation was found to impact the transcript abundance of all genes examined. Instead of the increased transcript abundance observed in the parent after treatment, when examining the *ΔoxsR* mutant: the *hvo_0040* and *hvo_1043* transcripts were detected at a constitutively low level throughout the time course, while the *hvo_0811* and *hvo_0037* transcripts were only modestly increased in abundance in the early stages of treatment. Counter to the other genes, the *hvo_0039* transcripts were found to be of greater abundance in the early stages of hypochlorite treatment in the *ΔoxsR* mutant compared to the parent. Overall, the qRT-PCR results suggest that after cells are exposed to hypochlorite, OxsR can act as a transcriptional activator (*e.g., hvo_0040*, *hvo_0337*, *hvo_0811* and *hvo_1043* regulation) as well as a repressor (*e.g., hvo_0039* regulation). Furthermore, the 3-fold increase in OxsR protein abundance during hypochlorite stress may in part be due to an increase in transcript level.

**Figure 4.**
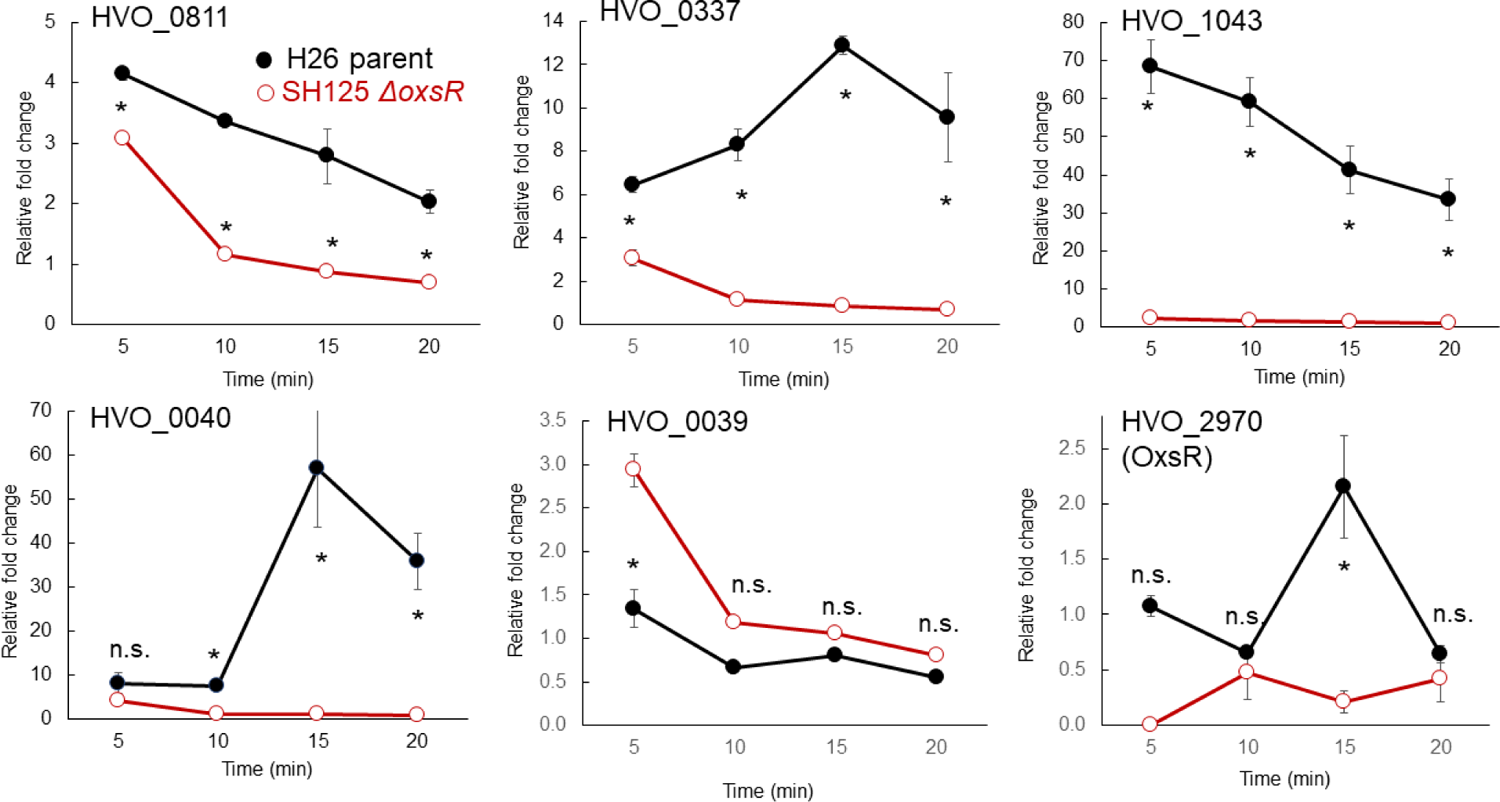
Influence of hypochlorite stress and *oxsR* on the expression of select genes identified by ChIP-seq analysis. Relative-fold change represents the transcript abundance ratio of NaOCl: mock treated cells. The *H. volcanii* H26 parent and SH125 (*ΔoxsR*) mutant cells were grown in GMM to exponential phase and treated with 0 and 2.5 mM NaOCl for 5, 10, 15 and 20 min. Total RNA was extracted and used for quantitative real-time (qRT) PCR. Levels of the gene expression were normalized to the internal reference *ribL* (HVO_1015, one-fold). Targets of qRT-PCR are indicated as gene locus tag numbers within each panel. *Significant differences between the parent and mutant by the student t-test analysis (p-value ≤ 0.05). n.s., not significant. All data are expressed as mean ± S.E.M. See methods for details.

### Conserved DNA motif in the 5’ region of a subset of genes identified in the OxsR ChIP-seq dataset

*De novo* motif analysis of the DNA sequences of the intergenic regions identified by ChIP-seq was performed. This analysis was executed to identify conserved DNA motifs in the OxsR-bound intergenic sites and to determine whether the location of these motifs relative to the core promoters correlated with expression change. Similar regions from related haloarchaeal genomes were included in the analysis to enhance DNA motif identification. DNA motifs predicted by MEME MAST (70) to be common to the datasets were subsequently used to scan the *H. volcanii* genome by FIMO (71). FIMO analysis was performed: i) to identify the location of the DNA motifs on the *H. volcanii* genome; and ii) to calculate the significance of the findings in terms of false discovery rate (*q*-value) and probability (*p*-value). The FIMO identified sites were then compared to the current ChIP-seq and previously published SILAC datasets, with the latter based on the intergenic regions 5’ of genes encoding proteins of differential abundance after hypochlorite treatment as detected by quantitative SILAC-based MS analysis (69). A semi-palindromic DNA motif with evenly spaced CG repeats, <u>CG</u>Gn<u>CG</u>nG<u>CG,</u> was identified within OxsR-bound regions (where n and underline represent the bases of low conservation and palindrome, respectively, E-value 2.5 x 10^-132^; **Figure 5A**). This motif was detected at 89 sites on the *H. volcanii* genome at a *p*-value < 0.00001; with about one-third of the sites corresponding to 5’ regions associated with the ChIP-seq and SILAC datasets. Of the top seven sites identified at a *q*-value < 0.05, six clustered to at least one of the two datasets (**Figure 5B**). Further analysis of the top sites revealed the palindromic DNA motif of CG-repeats was generally positioned 5’ of BRE/TATA box consensus sequences presumed to serve as RNA polymerase binding sites; as depicted for *hvo_1043, hvo_1342*, *hvo_1668*, and *hvo_0040* regions (**Figure S6A-D**). The exception was *hvo_0039*, which immediately 5’ of its start codon had a CG-rich region that was not closely related in sequence to the CG repeat but was conserved in diverse *Haloferax* species (**Figure S6D, upper**), which may explain the repressive function of OxsR on the transcript levels of this gene. Thus, the positioning of the CG-rich motifs in relation to the basal promoter element was consistent with the up- or down-regulation in transcript levels observed for these genes by qRT-PCR (previous section). However, the CG-rich motif did not fully explain all OxsR-dependent activities as: 1) not all intergenic regions bound by OxsR have this CG-repeat motif and 2) the 12-fold increase observed for *hvo_0337* (glutaredoxin) transcripts during hypochlorite stress requires *oxsR,* yet the gene does not encode the CG-rich motif in its promoter region. Nonetheless, these results suggest for a subset of genes that placement of a CG-rich repeat upstream (*e.g., hvo_0811*, *hvo_0040* and *hvo_1043*) or downstream (*e.g., hvo_0039*) of the BRE/TATA box promoter consensus element may facilitate the ability of OxsR activate or repress gene expression in the presence of hypochlorite, respectively.

**Figure 5.**
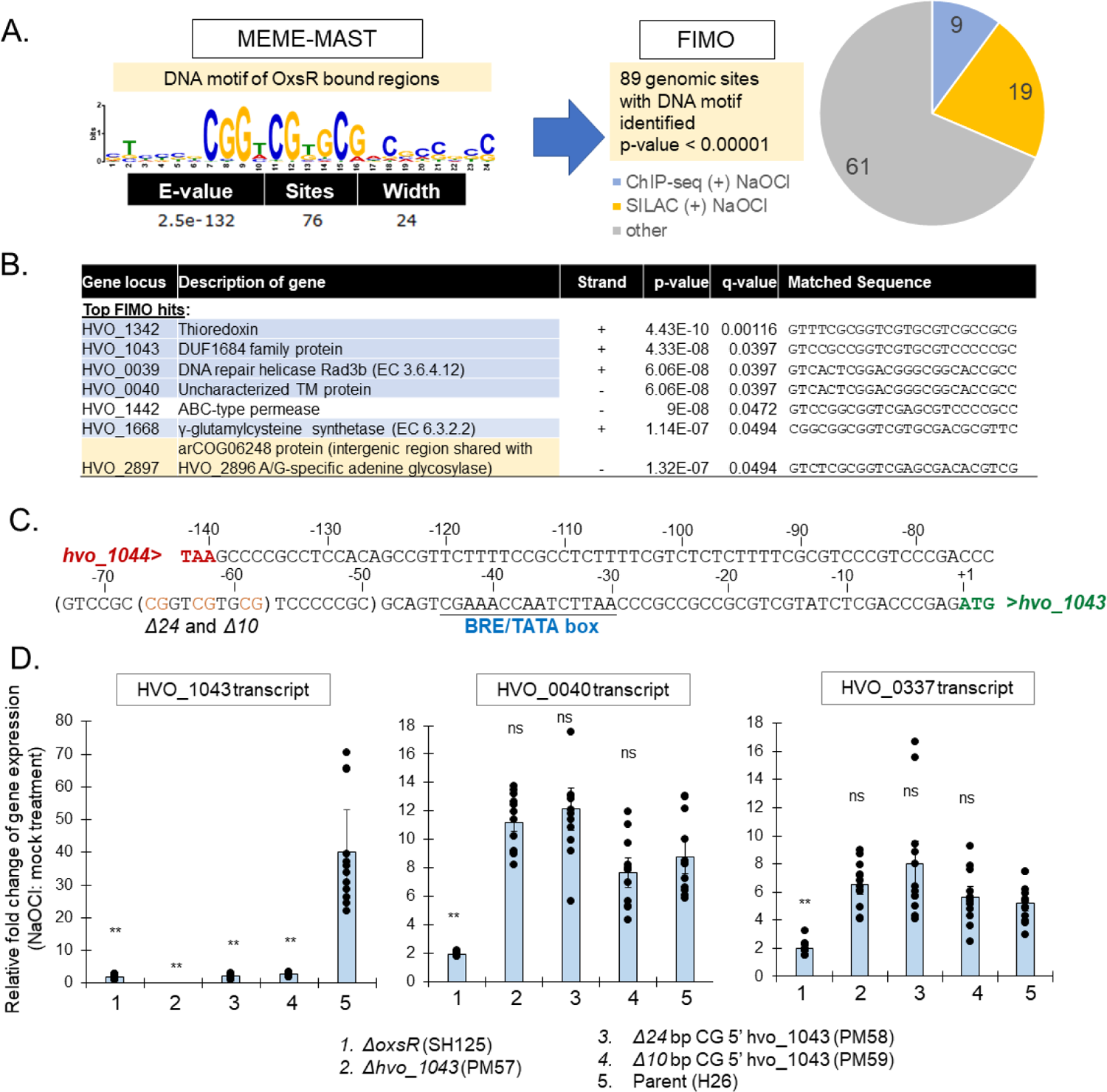
CG-rich DNA motif associated with OxsR-mediated transcriptional activation. A) CG-rich motif identified to be enriched in OxsR-bound ChIP-seq DNA sequences. The motif was identified by MEME-MAST analysis and was input into FIMO to scan the *H. volcanii* genome. The 89 sites identified by this approach at a *p*-value < 0.00001 were compared to the OxsR ChIP-seq (blue) and SILAC-based MS (orange) datasets. B) FIMO hits of the CG-rich motif detected at a *q*-value ˂ 0.05 with the sites common to the ChIP-seq (blue) and SILAC-based MS (orange) datasets highlighted. C) CG-rich DNA motif upstream of the BRE/TATA box of *hvo_1043* targeted for deletion. Parenthesis, CG-rich repeat region targeted for mutagenesis with the CG repeat in orange (Δ24 spans −50 to −74 and Δ10 spans −58 to −67). Red, stop codon of *hvo_1044*. Underlined, BRE/TATA box region. Green, start codon of *hvo_1043*. D. qRT-PCR analysis of deletions of CG-rich repeat 5’ of *hvo_1043*. *H. volcanii* strains were grown in GMM to early log phase (OD600 of 0.3 to 0.5) and exposed to 2.5 mM NaOCl or a mock control for 15 min. Transcript levels were monitored in the mutant strains as indicated. The internal reference *hvo_1015* normalized levels of gene expression were at one-fold relative fold change. Targets of qRT-PCR are indicated for each panel by gene locus tag number. Strains used for preparation of RNA are indicated on the x-axis. Significant differences between the parent and mutants by the student t-test analysis (**, p-value ≤ 0.01; n.s., not significant) (Exp./Bio: 4; Tech: 3 replicates). See methods for details.

### CG-rich motif required for the OxsR-dependent increase in *hvo_1043* transcript levels after exposure to HOCl

To determine if the CG-rich DNA motif was important for the observed OxsR-dependent increase in *hvo_1043* transcript abundance during hypochlorite stress, a mutagenesis approach was used. Strains with deletions in the CG-rich motif upstream of the BRE/TATA box consensus sequence of the *hvo_1043* operon were constructed and compared to the H26 parent and *ΔoxsR* mutant by qRT-PCR (**Figure 5C**). A *Δhvo_1043* coding sequence mutant was also constructed as a negative control. By this approach, a 40-fold increase in the abundance of *hvo_1043* transcripts was observed after 15 min of hypochlorite treatment in the H26 parent that was not apparent in the *ΔoxsR* or the CG-rich motif mutant strains (**Figure 5C**). The *hvo_1043* transcripts were detected only at basal levels in the *ΔoxsR* and CG-motif mutant strains. As expected, *hvo_1043* transcripts were undetectable in the *Δhvo_1043* mutant. As an added control, the transcripts of other genes (*hvo_0040* and *hvo_0337*) were confirmed to be unaffected by the CG-rich motif deletions that were specifically integrated upstream of *hvo_1043*. Based on these results, the CG-rich motif identified upstream of the BRE/TATA box consensus sequence of *hvo_1043* appears important for transcriptional activation of this operon by OxsR during hypochlorite stress. Because of the close sequence similarity of this GC-rich motif upstream of other OxsR-bound operons, we hypothesize that OxsR may bind this motif to regulate expression of at least a subset of genes within its regulon.

### Conserved residues and 3D-structural modeling of OxsR

As OxsR was associated with recovery from hypochlorite, which is a potent oxidant, the primary amino acid sequence of OxsR was inspected for conserved residues that may sense oxidant, such as Cys, His and Met (4). Multiple amino acid sequence alignment revealed that a cysteine (corresponding to OxsR C24) was highly conserved in the N-terminal region of OxsR homologs from diverse archaea (**Figure 6A**). OxsR was further analyzed by 3D-modeling to predict the location of C24 in the protein structure. The 3D-structure of OxsR residues 18-121 (104 of 123 aa total; 85%) was modeled at >99.8% confidence using the Phyre2 web portal (72). OxsR residues 1-123 were also modeled by RoseTTAfold (73). The alignment scores of the two OxsR 3D-models were at a mean square deviation (RMSD) of 3.47 Å and template modelling (TM)-score of 0.52, suggesting the models were generally related. The major exceptions were: i) residues 26-46 which were structurally undefined in the Phyre2-model and ii) residues 1-17 which were not modeled by Phyre2 and found in multiple configurations by RoseTTAfold. The highest scoring template by Phyre2 was the X-ray crystal structure of *Methanosarcina mazei* MM_1094 (PDB: 3R0A), an uncharacterized TrmB family member of arCOG02242, which has two intersubunit disulfide bonds formed between C6 and C17. The single cysteine (C24) of OxsR aligned with C17 of MM_1094. The OxsR quaternary structure was also predicted by comparison to two other high scoring (>99% confidence) templates including: an assembly of X-ray crystal structures of the *Sulfolobus acdiocaldarius* AbfR2 (Saci_1223) (PDB: 6CMV)(74) and *Streptococcus pneumoniae* FabT in complex with DNA (PDB: 6JBX)(75). The quaternary model generated by this approach suggested OxsR could form a homodimer linked by an intersubunit disulfide bridge at C24 that would join anti-parallel α helices of the two subunits (**Figure 6B**). A separate homodimer interface formed between two anti-parallel α helices at the C-terminus of OxsR was also predicted.

**Figure 6.**
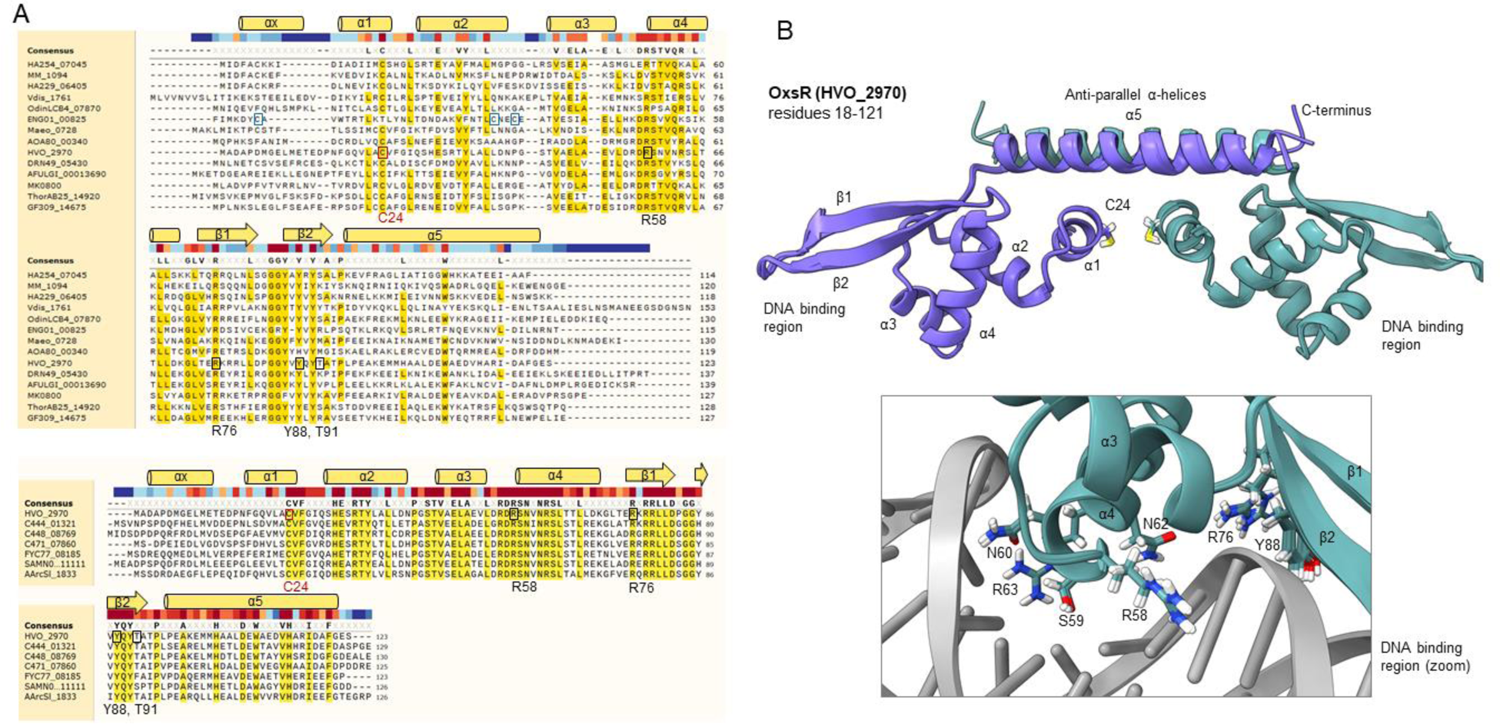
Conserved residues and 3D-structural model of OxsR. A) Multiple amino acid sequence alignment of OxsR (HVO_2970) to TrmB family proteins. Upper: OxsR aligned to representative homologs from diverse archaea. *Crenarchaeota* (Vdis_1761), *Thaumarchaeota* (DRN49_05430), *Thermoplasmata* (AOA80_00340), *Methanopyri* (MK0800), *Methanomicrobia* (MM_1094), *Methanococci* (Maeo_0728), *Archaeoglobi* (AFULGI_00013690), *Nanoarchaeota* (HA229_06405), *Aenigmarchaeota* (ENG01_00825), *Diapherotrites* (HA254_07045), *Thorarchaeota* (ThorAB25_14920), *Odinarchaeota* (OdinLCB4_07870) and *Lokiarchaeota* (GF309_14675). Lower: OxsR aligned to representative homologs from diverse families of haloarchaea. *Halorubraceae* (C471_07860 and AArcSl_1833), *Natrialbaceae* (FYC77_08185), *Haloarculaceae* (C444_01321), *Halococcaceae* (C448_08769), and *Halobacteriaceae* (SAMN05216226_111111). Residues of OxsR discussed in text indicated by black and red boxes and numbered below the alignment. Blue boxes indicate cysteine residues in the N-terminal region of *Aenigmarchaeota* homolog (ENG01_00825). Predicted α helices and β strands of OxsR indicated above the alignment. Residues at >50% and >95% amino acid sequence identity are indicated by yellow highlighting (upper and lower panels, respectively) with the bar colored dark red to dark blue above the sequence indicating residues of high to low conservation. B) 3D-structural model of OxsR (HVO_2970). Ribbon diagram of OxsR in homodimeric configuration (chain A and B in purple and cadet blue, respectively). Residue numbering and secondary structure indicated for chain A. The 3D-structural model generated by RoseTTAfold for residues 18-123 is displayed. Arrangement of the 3D-model into a homodimer was by comparison to the X-ray crystal structure of the biofilm regulator *Sulfolobus acidocaldarius* AbfR2 (Saci_1223; PDB: 6CMV). DNA interactions were predicted by comparison to the X-ray crystal structure of the *Streptococcus pneumoniae* FabT:DNA complex (PDB: 6JBX).

### Cysteine residue is important for OxsR function

To determine whether OxsR C24 is important for sensing oxidative stress, the *oxsR* genomic locus was modified by site-directed mutagenesis. A strain was constructed to express OxsR C24A with a C-terminal HA tag (OxsR-HA C24A) from the *oxsR* locus similarly to the OxsR-HA form. Using this strain, the role of the conserved C24 was monitored based on: i) the recovery of cells from hypochlorite stress and ii) the relative fold-change in *hvo_1043* transcript abundance after exposure of these cells to hypochlorite. The single amino acid exchange of OxsR from C24A reduced the ability of cells to recover from hypochlorite stress, while having little if any impact on growth in the presence of the mock control (**Figure 2AB**). When using *hvo_1043* transcripts to monitor OxsR C24 function, the *oxsR*-HA C24A integrant strain was found to be severely impaired in its ability to upregulate the abundance of *hvo_1043* transcripts in the presence of hypochlorite, to a level comparable to the *ΔoxsR* mutant (**Figure 7B**). By contrast, the parent (H26) and the OxsR-HA integrant strains displayed a robust increase in *hvo_1043* transcript levels when exposed to hypochlorite (**Figure 7B**). To determine whether the C24A had an impact on OxsR protein abundance in the cell, the levels of OxsR-HA and OxsR-HA C24A were compared by anti-HA tag immunoblotting analysis (**Figure 7C**). The anti-HA antibodies were found to be specific to the strains expressing OxsR-HA variants, as no signal was detected in the parent strain devoid of the HA tag. Furthermore, on visual inspection of the immunoblots, the OxsR-HA C24A and OxsR-HA were comparable in protein abundance, suggesting the C24A modification did not impact OxsR expression or stability. While amino acid exchange at C24 did not alter OxsR protein abundance, it did eliminate the ability of OxsR to facilitate the recovery of cells and upregulate the level of *hvo_1043* transcript during hypochlorite stress. Thus, C24 appears to be important for OxsR function as a transcriptional activator when strong oxidants of cellular thiols are introduced into the environment.

**Figure 7.**
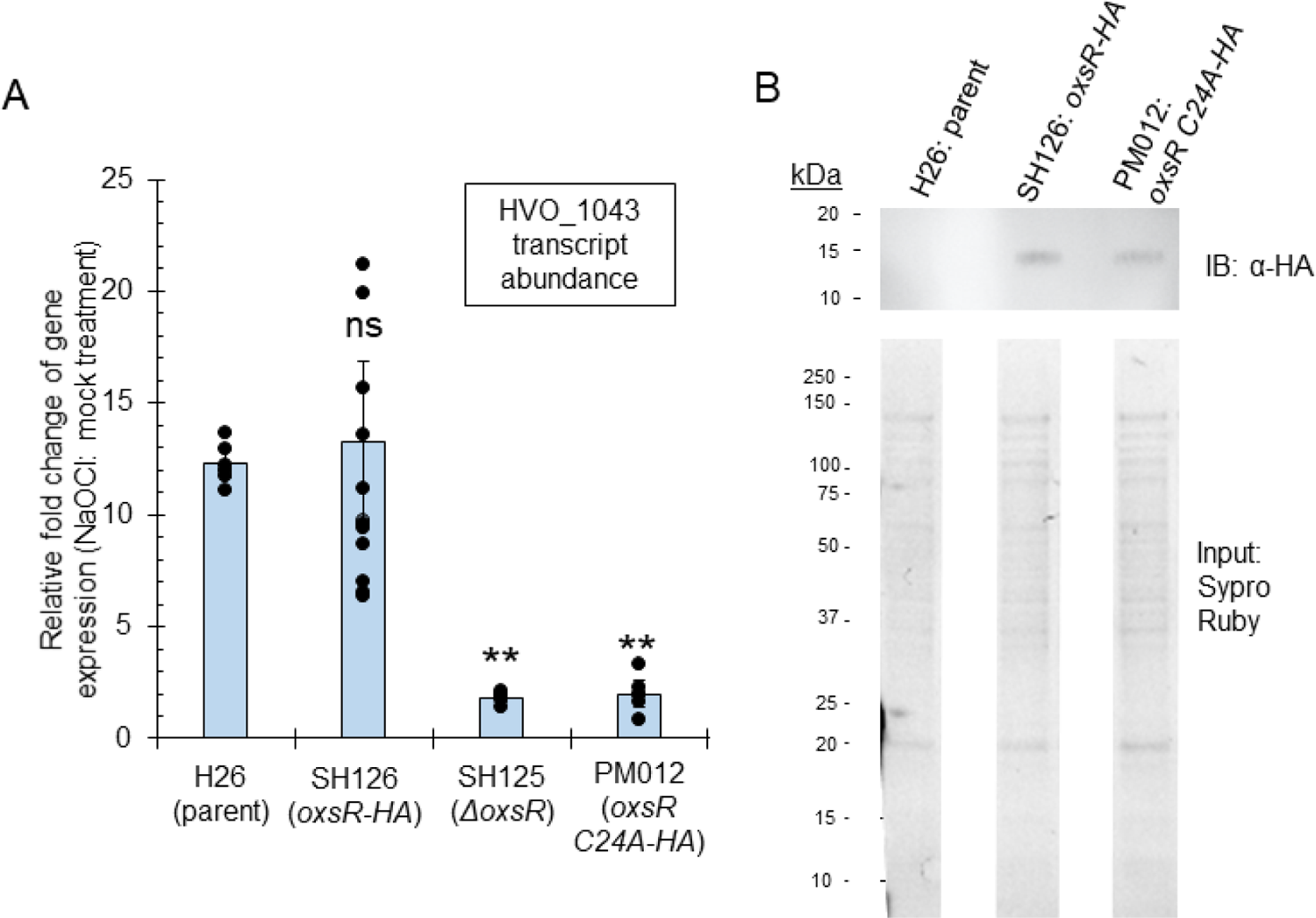
Conserved cysteine residue (C24) important for OxsR function. A) Relative-fold change of *hvo_1043* transcript levels during hypochlorite stress. *H. volcanii* strains as indicated on the x-axis were grown to early exponential phase in GMM and treated with 0 and 2.5 mM NaOCl for 15 min. This timeframe was found to result in a 10- to 40-fold increased abundance of *hvo_1043* transcripts in the parent strain. Total RNA was extracted and used for qRT-PCR analysis. The internal reference *hvo_1015* normalized levels of the gene expression were at one-fold relative fold change. Significant differences between the parent and mutant by the student t-test analysis (**, p-value ≤ 0.001; *, p-value ≤ 0.05). n.s., not significant. (Exp./Bio: 2-4; Tech: 3 replicates). B) Detection of OxsR-HA with and without the C24A variant in *H. volcanii* cells. Strains were grown to early log phase (OD600 of 0.3 to 0.5). Upper panel: Immunoprecipitates (10 µl per lane) were separated by reducing 15% SDS-PAGE for 2 h at 100 V. Proteins were detected by immunoblotting analysis using anti-HA tag HRP (# ab1190) antibodies at 1:20,000 dilution and ECL Prime. Signal was visualized after a 30 sec exposure. Molecular mass standards were Precision Plus Protein Kaleidoscope. Lower panel: Total protein input separated by reducing 12% SDS-PAGE prior to immunoprecipitation detected by Sypro Ruby staining is included as control. Samples were normalized as 4 µl (0.04 OD600 units cell pellet) per lane. See methods for details.

## Discussion

Here we advance knowledge of oxidative stress signaling in prokaryotes by associating TrmB-like single winged-helix DNA binding domain proteins from diverse archaea as thiol-based transcriptional regulators of oxidative stress response. TrmB is a large and diverse protein family that accounts for over 10% of the total TFs in archaea and 0.5% of the TFs in bacteria; however, thiol-based sensory mechanisms were not previously reported for this family. Using the TrmB-like OxsR of *Haloferax volcanii* as a model, we demonstrate that this protein functions as a transcriptional activator and repressor of a large gene co-expression network associated with oxidative stress response. A conserved cysteine residue serves as the thiol-based sensor for this function and likely forms an intersubunit disulfide bond during hypochlorite stress that stabilizes a homodimeric configuration of OxsR with enhanced DNA binding properties.

Our study relies upon *H. volcanii* cells grown in glycerol minimal medium and exposed to hypochlorite. Thus, the antioxidants present in complex, undefined medium did not complicate our experiments. The hypochlorite used for the environmental cue is a reactive species common to biological systems of high chloride concentration (57). Furthermore, the carbon and energy source, glycerol, was of relevance to hypersaline habitats, where algae accumulate large amounts of glycerol for osmotic stabilization, which is released into the environment (76) for use by heterotrophs such as *H. volcanii* which prefers glycerol over glucose (77).

Evidence suggests OxsR binds a conserved CG-rich DNA motif in promoters of select genes. Promoters with CG-rich motifs located 5’ of the TATA box and BRE consensus were activated by OxsR, whereas those 3’ were repressed. These data suggest that the motif may serve as an OxsR binding site and that the positioning of this site in relation to core promoters (TATA binding protein / transcription factor B binding sites) may determine whether OxsR functions as an activator or repressor (78, 79).

While the CG-rich DNA motif is common to many of the sites bound by OxsR, not all of the OxsR-bound sites share this motif. One possible explanation for this finding is that other protein factors could associate with OxsR and influence the type of binding site recognized by this TF. Consistent with this hypothesis, some gene induction at early time points following exposure to hypochlorite is still observed in the *ΔoxsR* strain for certain OxsR activated genes (*hvo_0811*), and late repression is possible for other genes (*hvo_0039*), invoking the involvement of another regulator. One candidate is HVO_1360, a small TrmB family protein that shares 28% amino acid identity with OxsR and similarly clusters to arCOG02242 (**Figure S7A**). 3D-structural modeling suggests HVO_1360 forms a wHTH domain flanked on each side by two α-helices much like OxsR (**Figure S7**). Thus, HVO_1360 could potentially form a heterodimer with OxsR that is stabilized by an intersubunit disulfide bond at the anti-parallel α interface of these two subunits (OxsR C24 bound to C21 or C15 of HVO_1360). Formation of this type of complex could alter the DNA sites bound by OxsR and present in the ChIP-seq dataset. An alternative explanation is that OxsR has an extensive DNA footprint with limited to no recognizable motif as seen for other TFs (discussed in detail below).

TFs with thiol-based redox switches are common in bacteria (4, 5). These TFs are generally classified into 1-Cys and 2-Cys based on sensing through one or two cysteine residues, respectively. Methionine or histidine residues, flavin cofactors, iron, iron-sulfur clusters, and heme centers are also used in bacterial TFs to sense redox status and can be found as added sensors in the thiol-based TFs. ROS or other redox-active compounds can cause specific modifications that lead to conformational changes of these TFs and result in the loss, gain or alteration of their DNA binding activity. Examples of bacterial TF thiol-based modifications include inter- or intra-subunit disulfide bond formation, S-thiolation (mixed disulfides of proteins and low molecular weight thiols), cysteine phosphorylation, and thiol-S-alkylation. These modifications can lead to transcriptional activation, repression or derepression depending on the TF. Many bacterial TFs with thiol-based switches are inactivated by redox stress leading to transcriptional derepression or deactivation, *e.g.,* 1-Cys OhrR (11, 12), 2-Cys OhrR (13), SarA/MgrA (14), PerR (15), HypR (16), YodB (17), QsrR (18), MosR (19), SarZ (20). One of the most versatile groups of thiol-based bacterial TFs activated by stress are the OxyR homologs of the LysR family including 2-Cys and 1-Cys type. In the presence of peroxide, nitric oxide (^·^NO) or oxidized glutathione (GSSG), OxyR forms intramolecular disulfide bonds (2-Cys OxyR) or other post-translational thiol modifications that can transform the TF into a transcriptional activator (21) or repressor (22) or lead to derepression (23). In the oxidized state, OxyR forms tetramers that bind DNA with an extensive footprint (∼50 bp) composed of repeating spaced elements of limited sequence similarity (80, 81). The well-known bacterial SoxRS operon is also activated by oxidative stress, with its DNA binding activity triggered by the oxidation or nitrosylation of [2Fe-2S] clusters (24, 25). By contrast, bacterial FNR homologs require the presence of an intact 4Fe-4S cluster (otherwise disrupted by oxygen) for function as global transcription regulator when oxygen becomes scarce (26).

Thiol-based redox switch TFs are also found in archaea and eukaryotes. Prior to our work, thiol-based redox switches were identified in TFs of archaea but limited to the ArsR family of transcriptional repressors MsvR (31–33) and SurR (27–30). Upon shifts to oxidizing conditions, these 2- and 5-Cys TFs form intra- and intersubunit disulfide bonds (SurR (29) and MsvR (32), respectively) that result in TF inactivation and transcriptional deactivation and/or derepression. In yeast, Yap1 is identified as a basic leucine zipper (bZIP) TF that uses a thiol-based redox switch to function as a central regulator of oxidative stress response pathways (82, 83). In the presence of H_2_O_2_, Yap1 forms intramolecular disulfide bonds that alter its conformation and mask its nuclear export signal. These changes promote the accumulation of Yap1 in the nucleus and stimulate transcriptional activation of its regulon (84), with formation of an intermolecular thiol intermediate between Yap1 and the thiol peroxidase Gpx3 inhibiting this activity (85). Thus, of the various domains of life, thiol-based TFs that are activated by oxidative stress and mediate a global response have yet to be reported in archaea.

Small wHTH domain proteins that cluster to arCOG02242 similarly to OxsR are characterized in *Crenarchaeota* including Lsr14, Smj12 and others (**Table S3**). Of these proteins, Lsr14 purifies as a homodimer and forms large footprints (> 50 bp) over its own promoter and the alcohol dehydrogenase (*adh*) promoter suggesting it functions as a transcriptional repressor (86–88).

The size of the footprints suggests that Lsr14 assembles from a homodimer into higher-order complexes in the promoter region (88). Lsr14 is also found to associate with other DNA binding proteins, such as the benzaldehyde-activated TF (Bald) and the chromatin remodeling proteins Sso7d and Sso10b (Alba), suggesting additional levels of regulation (88). While of low protein abundance in the cell, the related Smj12 displays activities which suggest it has a role in chromatin remodeling including binding DNA non-specifically, stabilizing the DNA double helix, and introducing positive supercoiling in DNA (89). More recently, Lrs14, AbfR1 and AbfR2 proteins of this arCOG group are correlated with biofilm formation, adhesion and motility (90, 91), with phosphorylation of AbfR1 Y84 and S87 found important for its binding to promoter regions (92).

Based on this study, we find OxsR to have common and distinct properties with the arCOG02242 group representatives that have been characterized. While a CG-rich motif appears important for the DNA binding and transcriptional activity of OxsR at promoter regions, the ChIP-seq enrichment peaks for OxsR were on average 700 bp (**Table S2**), which is wider than the peaks observed for other previously characterized halophilic TFs (40, 93). Furthermore, our ChIP-seq analysis revealed six operons to have multiple 5’ intergenic sites bound to OxsR (*i.e*., *hvo_A0618*, *hvo_2758*, *hvo_1875*, *hvo_0198*, *hvo_1043* and *hvo_1342*). These results suggest OxsR homodimers may form higher order structures and bind DNA with large footprints similarly to Lsr14. OxsR does not appear to be a highly abundant protein that non-specifically binds DNA, as specific sites were identified by ChIP-seq analysis, and OxsR-HA could not be detected by western blotting without prior enrichment by immunoprecipitation, which contrasts with the HA-tagged TF GlpR (94). Furthermore, unlike abundant proteins such as proteasomes, OxsR is detected in only four of the six whole proteome datasets reported in the Archaeal Proteome Project (ArcPP) (95). The most important distinction of OxsR with the previously characterized arCOG02242 members is that it is found to use a thiol-based sensor in the transcriptional response associated with recovery from hypochlorite stress.

Post-translational modification (PTM) appears important in regulating the activity of the arCOG02242 TFs. We find most members of this arCOG group have conserved cysteine residues located at predicted homodimer interfaces formed by either the antiparallel α1 and/or α5 helices (**Figure 6**, **Table S3**). These cysteine residues form intersubunit and intrasubunit disulfide bonds in the X-ray crystal structures of Sto12a and MM_1094 and, thus, could generally serve as redox sensors that influence TF homodimer stability. Ser/Thr and Tyr phosphorylation provides an additional type of post-translational regulation to consider in the signaling pathway of TFs of the TrmB family arCOG02242 group (92). While the Y84 and S87 phosphosites of AbfR1 are less conserved among the OxsR homologs than the OxsR C24, we find OxsR to have residues (Y88 and T91) analogous to the sites of AbfR1 phosphorylation suggesting PTM crosstalk between a thiol-switch and phosphorylation.

In this work we revealed the requirement of OxsR when *H. volcanii* is exposed to oxidative stress. This work nicely complements previous findings of RosR function in the haloarchaeal oxidative stress response (61–63), as the mode of DNA binding and mechanism of sensing oxidant appear quite distinct between OxsR and RosR (the latter yet to be determined). OxsR binding regions that were mapped by using a genome-wide ChIP-seq approach provide the biological roles not only as an activator for genes involved in amino acid and thiol transfer but a repressor for DNA repair system (**Figure 8**). The conserved DNA motif (CGGnCGnGCG) and cysteine residue are mainly contributed to the OxsR binding, however, other factors that interact with OxsR to find/bind to target DNA regions would be remained to be discovered. Overall, this work supports an emerging principle that OxsR which is widespread in most archaeal phyla plays a pivotal role against oxidative stress.

**Figure 8.**
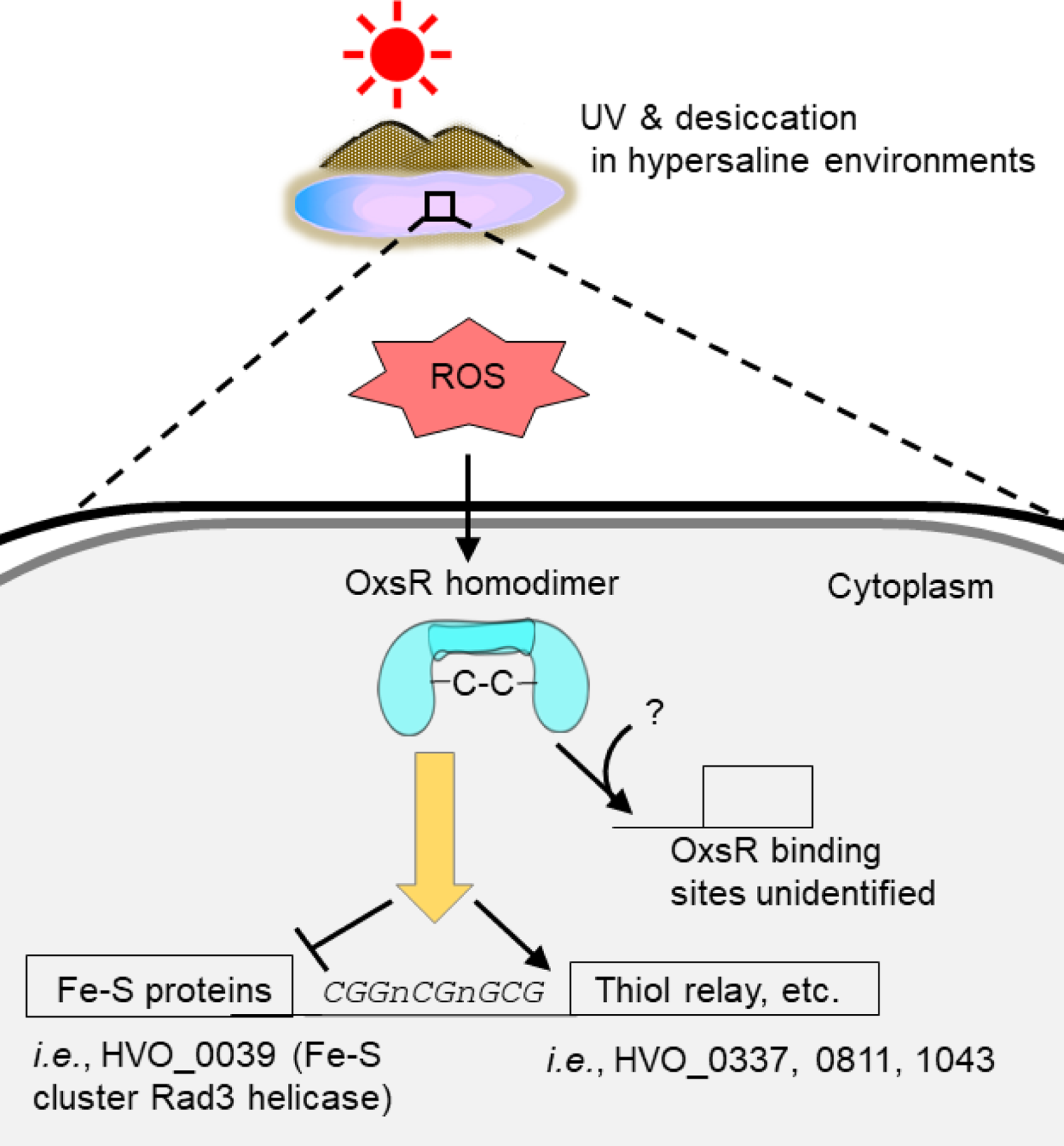
A proposed model of OxsR-mediated oxidative response in *H. volcanii*. The diagram shows that ROS generated from (a)biotic reactions is sensed by OxsR and then an intersubunit disulfide bond is formed at the conserved cysteine residue, followed by regulating gene expression (activation/repression and target DNA binding).

## Materials and Methods

### Materials

Biochemicals and analytical-grade inorganic chemicals were purchased from Fisher Scientific (Atlanta, GA), Bio-Rad (Hercules, CA) and Sigma-Aldrich (St. Louis, MO). Desalted oligonucleotides were from Integrated DNA Technologies (Coralville, IA). DNA polymerases and restriction enzymes were from New England Biolabs (Ipswich, MA) and Clontech Laboratories, Inc. (Mountain View, CA). Hi-Lo DNA standards were from Minnesota Molecular, Inc. (Minneapolis, MN).

### Strains, media and growth conditions

Details of strains, plasmids, and primer sequences used in this study are listed in **Table S4**. *E. coli* cultures were grown at 37 °C in Luria-Bertani (LB) medium. *H. volcanii* strains were grown at 42 °C in ATCC974 complex medium or glycerol minimal medium (GMM) as previously described (77). Liquid cultures were grown with rotary agitation at 200 rpm. Media was supplemented with 1.5% agar for plates and with novobiocin (Nv, 0.2 µg·ml^-1^) or ampicillin (Ap, 100 µg·ml^-1^) as needed. Manipulation of *H. volcanii* strains and DNA was as described by the *Halohandbook* (http://www.haloarchaea.com/resources/halohandbook/). Plasmids were generated in *E. coli* TOP10 and then transferred to *E. coli* GM2163 prior to transformation of *H. volcanii*. Strain and plasmid fidelity was confirmed by Sanger DNA sequencing (Eton Bioscience, Inc., San Diego, CA).

### Generation of *H. volcanii* mutant strains

The *H. volcanii ΔoxsR* mutant (SH125) was generated using *H. volcanii* H26 as the parent by a *pyrE2*-based pop-in/pop-out deletion method (48). The pre-deletion plasmid pJAM3380 was generated by ligation of a DNA fragment carrying *oxsR* and 5’ and 3’ flanking regions (about 600 bp each) into the HindIII to XbaI sites of plasmid pTA131. The *oxsR* region carried on pJAM3380 was generated by PCR using primer pair 1/2 and *H. volcanii* H26 genomic DNA as a template. Deletion plasmid pJAM3381 was constructed by inverse PCR using primer pair 3/4 and plasmid pJAM3380 as a template. Primer pairs used to screen for the SH125 mutant included 1/2, 3/4, and 5/6 (primers outside the deletion plasmid). Plasmid pJAM3388 was created by ligating a DNA fragment carrying *oxsR* into the NdeI and KpnI sites of pJAM809. The *oxsR* region carried on pJAM3388 was generated by PCR using primer pair 5/6 and *H. volcanii* H26 genomic DNA as a template. This plasmid and the empty vector (pJAM202c) were transformed into the *ΔoxsR* mutant (SH125) for complementation assay. A similar pop-in/pop-out strategy was used to generate the other *H. volcanii* strains including SH126, PM057, PM058, PM059, and PM012. Plasmid pJAM3389 used to integrate the *oxsR::HA* coding sequence into the *oxsR* locus was generated by inverse PCR using primer pair 7/8 and plasmid pJAM3380 a template. Strain SH126 was screened using primer pair 9/10. For site-directed mutagenesis to generate pJAM3901 (that encodes OxsR C24A with C-terminal HA tag), plasmid pJAM3389 and overlapping primer pair 11/12 were used to amplify a linear plasmid by PCR that was DpnI treated to remove template and ligated using a KLD Enzyme Mix (NEB). For deletion of *hvo_1043*, the *hvo_1043* region carried on pJAM3919 was generated by PCR using primer pair 27/28 and *H. volcanii* H26 genomic DNA as a template. Deletion plasmid pJAM3920 was constructed by inverse PCR using primer pair 29/30 and plasmid pJAM3919 as a template. Primer pairs used to screen for mutants included 27/28 (primers outside the deletion plasmid). The deletions of the CG-rich motif 5’ of *hvo_1043* were performed in a similar matter but used different primers. To generate the 24 bp deletion, primer pair 31/32 was used for inverse PCR with pJAM3919 as template. For the 12 bp deletion of the CG repeat, primer pair 33/34 was used for inverse PCR with pJAM3919 as template. All constructed plasmids were verified by the Sanger DNA sequencing method (Eton Bioscience, Research Triangle Park, NC). The SH125 (*ΔoxsR*) mutant was also confirmed by comparison to the H26 parent using the breseq pipeline (96) for whole genome sequencing (**Table S1**).

### Hypochlorite stress growth assay

For hypochlorite stress growth assays, the following method was used. The *H. volcanii* strains were streaked from −80 °C glycerol stocks onto glycerol minimal medium (GMM) plates and incubated for 5 days in a closed plastic zippered bag at 42 °C. Five isolated colonies were inoculated into 50 ml of GMM in 250 ml Erlenmeyer flasks. Cells were grown to late log phase (OD_600_ of 0.85-0.95) at 42 °C (200 rpm, rotary shaking). Cells were diluted with fresh GMM to a OD_600_ of 0.1 unit in 95 ml final volume. Aliquots (5 mL) of this cell suspension were transferred into 12 loosely capped 13 × 100 mm culture tubes per strain type and incubated for 10 h at 42 °C with aeration using a mini rotator (Glas-Col from Terre Haute in the USA) at a max percent speed setting of 50. Once cells reached log phase (OD600 of 0.4-0.6), half of the 12 tubes were randomly selected for supplementation with or without 5 mM NaOCl (sodium hypochlorite reagent grade, available chlorine 10-15%, Sigma-Aldrich, #425044-250mL), which forms hypochlorite in solution. The tubes were returned to the mini rotator and cell growth was monitored for 10 days at OD600 using a Spectronic 20+ spectrophotometer (ThermoSpectronic, Filter: 600-950nm). The experiment was determined to be reproducible (n = 12 total; 6 tubes per strain plus condition for each of 2 experiments). Initial analysis by circular rotary shaking yielded variable results, most likely due to the microaerobic conditions which promoted extensive incubation times after exposure to hypochlorite.

### Quantitative real-time reverse transcriptase PCR (qRT-PCR) analysis

*H. volcanii* strains for qRT-PCR analysis were streaked from −80°C glycerol stocks onto defined minimal media (GMM) and incubated for five days in a sealed plastic zippered bag at 42°C. Isolated colonies were inoculated into 20 ml of GMM in 125 ml Erlenmeyer flasks. Cells were grown to early log phase (OD_600_ of 0.3-0.5) at 42°C (200 rpm). For each strain, aliquots (1 ml) of the cell culture (20 ml) were transferred to 1.5 ml microcentrifuge tubes (RNase-free), and each sample was exposed to 2.5 mM of NaOCl for different times (5, 10, 15, and 20 min) at 42°C (200 rpm). A mock control was included for comparison of each time point. Total RNA was isolated from cells using TRI Reagent (#T9424) according to the supplier (Sigma-Aldrich). TURBO DNA-free™ Kit (AM1907) was used to remove contaminating DNA from the RNA samples according to the supplier’s recommendations (Invitrogen). Only RNA samples with DNA below the limit of PCR detection were further processed. RNA integrity was confirmed by mixing samples 1:1 in 2X RNA loading dye (B0363S; New England Biolabs) and separating by 0.8 % (w/v) agarose gel electrophoresis in 1X TBE. Only RNA (2 ng) samples with no apparent degradation served as the template for qRT-PCR in 20 μl reactions. The Luna Universal One-Step RT-qPCR kit (E3005L) was used for qRT-PCR analysis following the protocol described by the supplier (NEB) by using a CFX96 real-time C1000 thermal cycler (Bio-Rad). The reverse-transcription was performed under conditions of 55°C for 10 min. The quantitative real-time PCR was performed under conditions of 40 cycles at 95°C for 1 min, 95°C for 10 s, and 56°C for 15 s. An extension of 60°C for 30 s was performed followed by determination of the melting curve under conditions of 95°C for 10 s and increase in temperature from 60°C to 95°C for 5 s each. A single peak revealed by the melting curve indicated a single product. The internal standard HVO_1015 was used to normalize the target mRNA levels, based on finding its transcript levels were unperturbed by HOCl stress. Genomic DNA served as the template to test different primer pairs for PCR efficiency. Primers with PCR efficiency between 95 % and 105 % are listed in **Table S4**. For the analysis for qRT-PCR, fold-ratios of relative-fold change in gene expression were calculated using the 2^-ΔΔCt^ method (Livak & Schmittgen, 2001) For the statistical analysis, the student’s t-test was conducted to compare the mean of gene expression between the H26 parent and *ΔoxsR* mutant strain (SH125). For the qRT-PCR analysis of the *Δhvo_1043* mutant (PM057), strains with deletions of the OxsR binding motif identified 5’ of the promoter region of *hvo_1043* (PM058 and PM059), and the *oxsR*-HA C24A integrant strain (PM012), the same method was used as explained above, except no time-course was performed. The strains were exposed to 2.5 mM NaOCl for 15 min, and a mock control was included for each strain. All experiments were performed in biological duplicate or quadlets and technical triplicate.

### Preparation for ChIP-sequencing and data analysis

Four single colonies of SH126 (*oxsR-HA* integrant) and two colonies of H26 were inoculated in 5-mL GMM and grown aerobically at 42 °C to early stationary phase to synchronize growth phase (OD600nm, ∼1.0) with shaking (200 rpm). Cells were transferred to fresh 100-mL GMM, and 2.5 mM NaOCl was added for 20 min when cells reached log phase (OD600nm, 0.3∼0.5) for the oxidative stress group. A mock control was included for comparison. ChIP-seq samples were prepared as the previous method with modifications (97). Briefly, to crosslink, 37% formaldehyde was added to the culture at the final concentration of 1% and the cell culture was incubated on a rocking platform for 20 min at room temperature. A final concentration of 0.125 M glycine was added to stop the crosslinking reaction and the whole cells were washed three times with cold basal salts buffer followed by storage at −80°C until sonication. The cell pellet was thawed and resuspended in 800 μl lysis buffer (50 mM HEPES, 140 mM NaCl, 1 mM EDTA, 1% (v/v) Triton X-100, 0.1% (w/v) sodium deoxycholate, pH 7.5) containing protease inhibitor cocktail (Thermo Scientific) to shear DNA by Bioruptor 300 sonication system (Diagenode) with 15 cycles of 30 s on and 90 s off at high magnitude. The sheared DNA was monitored to confirm the genomic DNA was a smear between 200-800 bp with the highest concentration of fragments ∼500 bp by 1.2% (w/v) agarose gel electrophoresis. For immunoprecipitation, the sheared DNA was immediately incubated overnight with the complex anti-HA tag antibody (Abcam, ab9110) and Dynabeads protein A (Invitrogen) at 4°C. Enriched DNA/OxsR complexes were eluted by adding 50 µl elution buffer (50 mM Tris, 10 mM EDTA, 1% SDS (w/v), pH 8.0) and incubation at 65°C for 10 min. Reverse crosslinking was performed by incubating in TE/SDS (10 mM Tris, 1 mM EDTA, 1% SDS) overnight at 65°C. DNA, of which the RNA was removed, was subsequently extracted by a phenol-chloroform method. Library preparation and deep sequencing were carried out at Duke sequencing core. Sequencing reads were trimmed and controlled their quality (Phred score > 30) by TrimGalore! wrapper pipeline (https://github.com/FelixKrueger/TrimGalore) with default parameters. The preprocessed reads were mapped (alignment rate > 95%) using Bowtie2 (98) to the *H. volcanii* DS2 reference genome (https://www.ncbi.nlm.nih.gov/genome/1149?genome_assembly_id=170797). The mapped reads were then sorted and indexed by Samtools (99). Peaks (cutoff, Qval < 0.05) were called by MOSAiCS (100) and checked for quality by ChIPQC (101). DiffBind was used to identify significant peaks that were present at least three of four biological replicates (102), and ChIPseeker was used to annotate the peaks (see the peak information in **Table S2**) (103). Peak heights reported represent the mean of the ratio of read counts in the IP sample vs input control. Integrative genomics viewer (IGV) was used for the manual evaluation of peak height and peak location, as well as for the data visualization (104). For all gene lists harboring peaks in their upstream coding region, a functional enrichment was performed with arCOG categories (67) based on the hypergeometric distribution test as described previously (105, 106). Code associated with this ChIP-seq analysis are freely available at https://github.com/amyschmid/OxsR_ChIP_WGS.

### Immunoprecipitation of OxsR-HA with and without the C24A mutation

*Hfx. volcanii* strains H26, SH126 (H26 *oxsR*:HA integrant) and PM012 (H26 *oxsR*:HA C24A integrant) were inoculated from ATCC 974 plates to glycerol minimal medium (GMM) (3 ml in 13 × 100 mm tubes) and grown to late log phase (OD600, 0.8-1.0). Cells were diluted to an OD600 of 0.03 into fresh GMM (25 ml in 250 ml Erlenmeyer flask) and grown to log phase (OD600, 0.8-1.0).

Cells were transferred to fresh GMM (200 ml cultures in 1 L Erlenmeyer flask) and grown to stationary phase (OD600, 1.6). Cells were harvested by centrifugation (10,000 × *g* for 30 min, 4 ᵒC). Cell pellets were washed with 150 ml 20 % salt water (SW, where 20 % is composed of 2.46 M NaCl, 88 mM MgCl_2_, 142 mM MgSO_4_, 56.3 mM KCl, 42 mM Tris-Cl, pH 7.5) by centrifugation (10,000 × *g* for 20 min, 4 ᵒC). Cell pellets were stored at −80 ᵒC for 2 days before used. Cell pellets were resuspended in 0.8 ml of 0.2 % (w/v) SDS in lysis buffer composed of 50 mM HEPES, 2 mM EDTA, 150 mM NaCl, 1 % (v/v) Triton X-100, 0.1 % (w/v) sodium deoxycholate, pH 8.0. Throughout the immunoprecipitation experiment, the buffers were maintained on ice and supplemented with protease inhibitor cocktail according to supplier (Sigma). The resuspended cells were incubated on ice for 15 min and subsequently sonicated with an aspiration probe for 10 cycles (10 pulses, 0.5 s on, 0.5 s off at 30 % amplitude) (Sonic Dismembrator Model 500 fitted with a Branson model 102C aspiration probe, Fisher Scientific and Branson Ultrasonics, Danbury, CT) with ice-slurry incubations of at least 1 min between cycles. The sonicated samples were centrifuged (16873 × *g* for 30 min, 4 ᵒC). The supernatant was transferred to a 15 ml Falcon tube (Fisher Scientific), supplemented with 1 µg of α-HA-antibody (ChIP Grade, product # ab9110, Abcam, Cambridge, MA) in 3 ml ice-cold ChIP lysis buffer and incubated for 4 h at 4 ᵒC with rocking. During this time, Protein A Dynabeads (50 µl, Invitrogen) were prewashed two times in a 2 ml microcentrifuge tube (natural color) with 1 ml 1 × phosphate buffered saline (PBS) at pH 7.4 (containing 8.0 g NaCl, 0.2 g KCl, 1.44 g Na_2_HPO_4_, and 0.24 g KH_2_PO_4_ per liter) (where the beads were washed by application of slurry to a magnet and removal of supernatant by aspiration). Beads were blocked by addition of 400 µl BSA in PBS buffer (1 mg·ml^-1^). The BSA-bead slurry was added to the pre-incubated sample and further incubated overnight (rocking at 4 ᵒC). After incubation, the supernatant was removed from the beads (via magnet and aspiration), and the beads were resuspended in ice-cold lysis buffer (1 ml). The bead-slurry was transferred to a fresh 2 ml tube, and the supernatant was removed by application to a magnet and aspiration. This wash step was repeated for a total of two times and followed by subsequent washing steps that were each repeated twice with the following buffers: wash buffer 1 (lysis buffer with 150 mM NaCl), wash buffer 2 [10 mM Tris-Cl, 2 mM EDTA, 25 mM LiCl, 1 % (w/v) Nonidet P-40, 1 % (w/v) sodium deoxycholate, pH 8.0] and TE buffer (10 mM Tris-Cl, 1 mM EDTA, pH 8.0) respectively. After washing, the beads were resuspended in 100 µl 2x SDS reducing buffer (125 mM Tris-HCl, pH 6.8, 20% glycerol, 4% SDS, 0.1% bromophenol blue and 5% β-mercaptoethanol) and boiled for 10 min. The samples were centrifuged (5,000 × *g* for 5 min, room temperature). The supernatant was stored in a fresh 2 ml microcentrifuge tube and analyzed by immunoblotting. The remaining supernatant was stored at −20 °C for future use.

### Immunoblotting (western) analysis

Immunoprecipitants (10 μl per lane) were separated by reducing 15% SDS-PAGE. Proteins were transferred to PVDF membrane at 4°C for 14.5 h at 30 V using the mini trans-blot module in transblot buffer (10 mM MES buffer pH 6 and 10% (v/v) methanol) according to the supplier’s instructions (Bio-Rad). The membrane was removed from the cassette, and the location of the gel and protein standards were marked on the membrane using a pencil. The membrane was placed upright in an 18 by 10 cm plastic container and rinsed with 30 ml of 1x TBST (50 mM Tris-Cl, pH 7.5, 150 mM NaCl, and 0.5 ml/L Tween 20) for 30 min. The membrane was soaked with 100% methanol and dried for 1 h at room temperature under laminar air flow. The membrane was reactivated with 100% methanol, washed briefly with 1X TBST three times, and blocked for 3 h at 4°C in 60 ml blocking buffer composed of TBST buffer supplemented with 5% (w/v) BSA (Sigma Life Science). During the blocking stage, the membrane was gently rocked using the Lab-Line Rocker on medium (5 or 6) setting. The blocking solution was replaced with a solution of anti-HA tag horseradish peroxidase (HRP) antibody (ab1190) diluted to 1:20,000 in 60 ml of the blocking buffer. The membrane was incubated with gentle rocking for 1 h at 4°C. The membrane was rinsed for 1.1 h at room temperature with 50 ml TBST buffer seven times using the high setting of the rocker. The HRP signal of the antibody: protein complexes was visualized on the PVDF membrane by chemiluminescence using 2 ml of Amersham ECL Prime from CGE Healthcare Life Sciences and exposure to the iBright FL1500 Imaging System (A44241).

### Computational prediction of OxsR-binding DNA motifs

DNA fragments identified by ChIP-seq to be bound to OxsR were tested for common DNA motif(s) using the MEME Suite v. 4.12.0 (70). Sequences were input into the *de novo* motif detection mode of MEME-MAST with the following parameters: any number of repeats, max width of 24 bp, and 3 output motifs. Two DNA sequence sets were used as input. The first set, which did not generate DNA motifs of high significance, included all DNA sequences bound by OxsR. The second set, which identified the CG-repeating DNA motif <u>CG</u>Gn<u>CG</u>nG<u>CG (</u>E-value reported in the text represents the expected number of sequences in a random database of the same size that would match the motif), was a subset of the OxsR-bound DNA sequences and was supplemented with analogous intergenic regions from related haloarchaeal genomes. These related regions were retrieved by comparing the deduced protein sequences of the flanking genes by Basic Local Alignment Search Tool using BlastP (protein-protein BLAST) (107). The DNA sequences 5’ of the genes encoding these homologs were retrieved using the graphics tool within NCBI nucleotide portal (https://www.ncbi.nlm.nih.gov/nuccore/). DNA motifs identified in this manner were compared to shuffled sequences to determine significance. DNA motifs found to be significant (based on uniqueness to the unshuffled sequences) were input into the FIMO algorithm (71) to scan the *Hfx. volcanii* genome database (db/upstream_Prokaryotic_Haloferax_volcanii_DS2_uid46845_2018-06-11.fna) of 4073 sequences and 1,428,983 residues. The DNA sequences used to identify the <u>CG</u>Gn<u>CG</u>nG<u>CG</u> motif, and the FIMO output of the genome scanning are provided in **Table S2**.

### Computational prediction of OxsR 3D-structure

Protein sequences from UniProtKB accession numbers D4GY41 and D4GXQ1 were used to model 3D-structures of OxsR and HVO_1360, respectively. The Phyre2 web portal (72) was used to predict structure by fold recognition threading. RoseTTAfold (73) was used for *de novo* structure prediction. The 3D-models were visualized and compared using ChimeraX (108).

### Data availability

The ChIP-seq data discussed in this publication have been deposited in NCBI’s Gene Expression Omnibus (109, 110) and are accessible through GEO Series accession number GSE196894 (https://www.ncbi.nlm.nih.gov/geo/query/acc.cgi?acc= GSE196894). The whole genomic sequence (WGS) data for the *ΔoxsR* mutant (SH125) and *oxsR-HA* integrant (SH126) are deposited with accession number PRJNA806939 in the Sequence Read Archive (SRA)(111). Code and input data associated with the ChIP-seq analysis are freely available for download via GitHub: https://github.com/amyschmid/OxsR_ChIP_WGS.

## Acknowledgments

Funds awarded to JM and AS to develop systems biology tools were through the Bilateral NSF/BIO-BBSRC program (NSF 1642283). Funds awarded to JM to determine redox regulation in archaea were through the U.S. Department of Energy, Office of Basic Energy Sciences, Division of Chemical Sciences, Geosciences and Biosciences, Physical Biosciences Program (DOE DE-FG02-05ER15650). Funds awarded to JM to provide evolutionary insight in biological systems were through the National Institutes of Health (NIH 5R01GM057498). Funds to AKS to understand mechanisms of stress response affecting cell growth and gene network evolution were provided by the National Science Foundation (NSF 1651117 and 1936024).

## Author contributions

SH, PM, AS and JM designed the research experiments, analyzed the data, and wrote the paper. PM, SH and RO generated the genetic constructs unique to this study. PM and LK performed the qRT-PCR analysis. SH, PM, AS, and JM analyzed the global datasets. AN, EW, and RC provided insight into OxsR disulfide bond formation. All authors performed research experiments associated with this paper and have read and approved of the paper. The authors do not have a conflict of interest to declare.

## Notes

https://github.com/amyschmid/OxsR_ChIP_WGS

https://www.ncbi.nlm.nih.gov/geo/query/acc.cgi?acc=GSE196894

### Competing Interest Statement

The authors have declared no competing interest.

