## Supplemental_Tables-S3-S4-FigS1-S7 for "TrmB family transcription factor as a thiol-based regulator of oxidative stress response"

**Running title:** TrmB-like thiol-based transcription factor

**Table of Contents**

**Table S1.** The breseq resequencing analysis of H26 and Δ*oxsR* mutant strains (*see* **Excel file, Table-S1.xlsx**).

**Table S2.** ChIP-seq, MEME-MAST, and FIMO datasets and associated analyses (*see* **Excel file, Table-S2.xlsx**).

**Table S3.** Representative TrmB family (PF01978) proteins of arCOG02242.

**Table S4.** List of strains, plasmids, and primers used in this study.

**Figure S1.** *H. volcanii* mutant strains generated in this study including SH125, SH126, PM012, PM057 and PM058, PM059.

**Figure S2.** Further analysis of the Δ*oxsR* mutant.

**Figure S3.** Peak loci in chromosome and plasmids by ChIP-seq analysis in the absence (left) or presence (right) of oxidative stress.

**Figure S4.** Functional enrichment of gene homologs associated with sulfur relay systems by STRING 11.5 analysis.

**Figure S5.** DUF1684 family protein HVO_1043 and its conserved $CX_7C$ motif suggest a role in thiol chemistry.

**Figure S6**. DNA motif common to OxsR bound regions and its location in relationship to the promoter consensus sequence element in the 5’ region of select gene homologs identified by ChIP-seq analysis.

**Figure S7.** TrmB family proteins HVO_1360 and OxsR (HVO_2970) are structurally related.

**Table S3.** Representative TrmB family (PF01978) proteins of arCOG02242[1].

| Protein | Organism | pI/Mr (kDa) | CC | Cysteine(s) | Phosphosites | Description(s) | Structure | Refs. |
|---|---|---|---|---|---|---|---|---|
| OxsR (HVO_2970) | *H. volcanii* | 4.46/ 13.7 | none | C24 required for activity; intersubunit (C24-C24) disulfide bond predicted | Y88, T91 conserved | mutant impaired in growth in the presence of hypochlorite; protein and transcript abundance upregulated by hypochlorite; transcriptional repressor/activator of genes associated with redox stress | 3D-model | This study |
| HVO_1360 | *H. volcanii* | 4.57/ 15.3 | α1, α4, α5 | C15, C21, C32 | Not conserved | Other *H. volcanii* member of this arCOG group | 3D-model | This study |
| MM_1094 | *Methanosarcina mazei* | 7.71/ 14.0 | none | intersubunit (C6-C17) disulfide bonds | Y83 conserved | Homodimer; subunit configuration appears altered in X-ray crystal structure | PDB: 3R0A | Protein Data Bank |
| Smj12 (SSO0458) | *Saccharolobus solfataricus* | 9.35/ 12.9 | α5 | C105 | Not conserved | Non-specific DNA binding; stabilizes double helix; introduces positive supercoiling; not abundant | | (1) |
| Ss-Lrs14 (SSO1108) | *Saccharolobus solfataricus* | 7.72/ 14.1 | none | C25, C110 | Y89 conserved | Homodimer; autorepressor; binds *adh* (alcohol dehydrogenase) promoter; generally displays large footprint; accumulates in late growth phases | | (2-4) |
| AbfR2 (Saci_1223) | *Sulfolobus acdiocaldarius* | 8.69/ 14.6 | α1, α4, α5 | None | Y88 conserved | Mutant impaired in biofilm formation; upregulated at transcript level during biofilm | PDB: 6CMV | (5-7) |
| Sa-Lrs14 (Saci_1242) | *Sulfolobus acdiocaldarius* | 8.64/ 13.5 | none | C15, C16, C26, C100 | Y81, T84 conserved | Mutant impaired in biofilm formation; upregulated at transcript level during biofilm | | (6, 7) |
| AbfR1 (Saci_0446) | *Sulfolobus acidocaldarius* | 8.91/ 13.2 | α5 | C105 | Demonstrated Y84(p), S87(p); phosphorylation important for DNA binding | Mutant: non-motile, increased extracellular polymeric substance (EPS) production, robust biofilm structure, upregulation of adhesive pili (*aap*), downregulation of archaellum (*fla*); AbfR1 binds *aap* and *fla* promoter regions in vitro; role as transcriptional | | (6, 8) |

| | | | | | | | | |
|---|---|---|---|---|---|---|---|---|
| | | | | | | activator and repressor; upregulated at transcript level during biofilm | | |
| Sto12a (ST1889) | *Sulfurisphaera tokodaii* | 9.1/ 12.5 | α5 | intersubunit (C15-C15) intrasubunit (C16-C100) disulfide bonds | Y81, S84 conserved | Homodimer; α5 antiparallel coiled-coil homodimer interface | PDB: 2D1H | (9) |

[1]CC, coiled-coil region predicted by DeepCoil (10). *Sulfolobus acidocaldarius* arCOG02242 members: Saci_0133, Saci_0102, Saci_1242 (Lrs14), Saci_1223 (AbfR2), Saci_0446 (AbfR1), Saci_1219.

**Table S3 References**

**Table S4.** List of strains, plasmids, and primers used in this study.

| **Strain, plasmid or primer** | **Description**[a] | **Source or Ref.** |
|---|---|---|
| **Strains:** | | |
| *E. coli* | | |
| TOP10 | $F^-$ *mcrA Δ*(*mrr-hsdRMS-mcrBC*) Φ80*lacZΔ*M15 *ΔlacX74 recA1 araD139 Δ*(*ara leu*) 7697 *galU galK rpsL* ($Str^r$) *endA1 nupG λ-* | Invitrogen |
| GM2163 | $F^-$ *ara-14 leuB6 fhuA31 lacY1 tsx78 glnV44 galK2 galT22 mcrA dcm-6 hisG4 rfbD1 rpsL136 dam13*::Tn*9 xylA5 mtl-1 thi-1 mcrB1 hsdR2* | New England Biolabs |
| *H. volcanii* | | |
| DS70 | wild-type isolate DS2 cured of plasmid pHV2 | (1) |
| H26 | DS70 *ΔpyrE2* | (2) |
| SH125 | H26 ΔoxsR | This study |
| SH126 | H26 *oxsR*:*HA* integrant | This study |
| PM012 | H26 *oxsR C24A:HA* integrant | This study |
| PM057 | H26 *Δhvo_1043* | This study |
| PM058 | H26 Δ24 bp 5'-GTCCGCCGGTCGTGCGTCCCCCGC-3' CG-rich motif 5' of the BRE/TATA consensus sequence of *hvo_1043* | This study |
| PM059 | H26 Δ10 bp 5' CGGTCGTGCG-3' CG-rich motif 5' of the BRE/TATA consensus sequence of *hvo_1043* | This study |
| **Plasmids:** | | |
| pTA131 | $Ap^r$; pBluescript II containing P*fdx-pyrE2* | (2) |
| pJAM809 | $Ap^r$ $Nv^r$; pJAM202c containing $P2_{rrn}$-*hvo1862-strepII* (*KpnI* site inserted upstream of StrepII coding sequence) | (3) |
| pJAM202c | $Ap^r$; $Nv^r$; pJAM202-derived control plasmid | (4) |
| pJAM3380 | $Ap^r$; pTA131 carries *oxsR* and ~700 bp flanking sequence (pre-deletion plasmid) | This study |
| pJAM3381 | $Ap^r$; pJAM3380 with *ΔoxsR* (deletion plasmid) | This study |
| pJAM3388 | $Ap^r$ $Nv^r$; pJAM809 with *oxsR* complementation plasmid | This study |
| pJAM3389 | $Ap^r$; pJAM3380 with *oxsR*-HA (integrant plasmid) | This study |
| pJAM3901 | $Ap^r$, pJAM3389 with *oxsR*-HA C24A (integrant plasmid) | This study |
| pJAM3919 | $Ap^r$; pTA131 carries *hvo_1043* and ~500 bp flanking sequence (pre-deletion plasmid) | This study |
| pJAM3920 | $Ap^r$; pJAM3919 with *Δhvo_1043* (deletion plasmid) | This study |
| pJAM3921 | $Ap^r$; pJAM3919 with ΔDNA binding motif 5' of *hvo_1043* (deletion plasmid) | This study |
| pJAM3922 | $Ap^r$; pJAM3919 with ΔDNA binding motif CG repeat (deletion plasmid) | This study |
| **Primers:** | | |
| 1. preKO_HVO_2970_HindIII_F | 5' ATTACAAGCTTCTTCGACAACGAACTCGTGA 3' | This study |

| | | |
|---|---|---|
| 2. preKO_HVO_2970_XbaI_R | 5’ TAGTTTCTAGAGTAGCTGCCGTAGTCCTCGT 3’ | This study |
| 3. KO_HVO_2970_FW | 5’ GCTCGCGGCCGACCG 3’; deletion HVO_2970 | This study |
| 4. KO_HVO_2970_RV | 5’ GCACACCTGTTCGCCGTG 3’; deletion HVO_2970 | This study |
| 5. CompleHVO_2970_NdeI_F | 5’ GTTACATATGGCCGACGCACCGGACATG 3’; complement HVO_2970 | This study |
| 6. CompleHVO_2970_KpnI_R | 5’ GATTAGGTACCCTACGACTCGCCGAAGGCGT 3’; complement HVO_2970 | This study |
| 7. TrmBLHAtag_F | 5’ CCCGGACTACGCCTAGGCTCGCGGCCGACCG 3’; integrate HVO_2970 HA tag | This study |
| 8. TrmBLHAtag_R | 5’ ACGTCGTACGGGTACGACTCGCCGAAGGCGTCG 3’; integrate HVO_2970 HA tag | This study |
| 9. ext_HVO_2970_FW | 5' GGCTCCGTACTACTTCGACA 3’ | This study |
| 10. ext_HVO_2970_RV | 5' TCTCGATAGCTTCGACCATC 3’ | This study |
| 11. HAoxsR_C24A3FW | 5’ ACAGGTCCTCGCGGCGGTCTTCGGCATCC 3’ | |
| 12. HAoxsR_C24A3RV | 5’ GGATGCCGAAGACCGCCGCGAGGACCTGT 3’ | This study |
| 13. Qset2 HVO_2970_F | 5’ AACTTCGGACAGGTCCTC 3’; qRT PCR HVO_2970 (*oxsR*) | This study |
| 14. Qset2 HVO_2970_R | 5’ TTGTCGAGTAGTGCGAGATA 3’; qRT PCR HVO_2970 (*oxsR*) | This study |
| 15. ribL qPCR FW1 | 5’-GCGAGTACATCACGGGTATC-3’ | This study |
| 16. ribL qPCR RV1 | 5’ CACTTCCTCTTCGACCTTCAG 3’ | This study |
| 17. HVO_0040 Primer pair 6_F | 5’ GTCGTCATGGGAGCGATGAT 3’ | This study |
| 18. HVO_0040 Primer pair 6_R | 5’ GCGACGTGGATTGTGAAGC 3’ | This study |
| 19. HVO_0039 Primer pair 3_F | 5’ GAGTTGGCGGAGTTGAAGGA 3’ | This study |
| 20. HVO_0039 Primer pair 3_R | 5’ TCGAAAATCTCATCGGGCGT 3’ | This study |
| 21. HVO_0811 Primer pair 2_F | 5’ CCTCTCTTCGATGTGCACCC 3’ | This study |
| 22. HVO_0811 Primer pair 2_R | 5’ GGATTGGTGGCAAAGAACCG 3’ | This study |
| 23. HVO_0337 Primer pair 2_F | 5’ CGCGAAGGTGAAGGACAAAC 3’ | This study |
| 24. HVO_0337 Primer pair 2_R | 5’ GTCTGGCCGCTTACCTCTTC 3’ | This study |
| 25. HVO_1043 Primer pair 1_F | 5’ GAAGGACCGCTATCTCGCTG 3’ | This study |
| 26. HVO_1043 Primer pair 1_R | 5’ GTCGTCGTATTCCCGGAGTT 3’ | This study |

| | | |
|---|---|---|
| 27. preKO1043HindIII500up | 5' TACAAGCTTCGGCGAGTCCTGCTGGTTCGAG 3' | This study |
| 28. preKO_HVO_1043_XbaI-507down | 5' ATTTCTAGACGTTCGCCCACCTCGATCACCTCGTCG 3' | This study |
| 29. KO_HVO_1043_FW | 5' CCGAAGCCGAGAACTGAGAGACGCG 3' | This study |
| 30. KO_HVO_1043_RV | 5' CTCGGGTCGAGATACGACGCGGCGG 3 | This study |
| 31. KO_OxsRMotif1043_FW | 5' GCAGTCGAAACCAATCTTAACCC 3' | This study |
| 32. KO_OxsRMotif1043_RV | 5' GGGTCGGGACGGGACGCGAAAAG 3' | This study |
| 33. KO_OxsRMotif1043_F2 | 5' TCCCCCGCGCAGTCGAAACCAATC 3' | This study |
| 34. KO_OxsRMotif1043_R2 | 5' GCGGACGGGTCGGGACGGGACG 3' | This study |

[a]$Ap^r$, ampicillin resistance; $Nv^r$, novobiocin resistance; $Str^r$, streptomycin resistance. $P2_{rrnA}$, rRNA promoter used for gene expression; *oxsR*, TrmB-like HVO_2970; HA, C-terminal hemagglutinin derived epitope tag., *ribL* (internal standard); HVO_2970, *trmBL* renamed *oxsR*. Underlined nucleotides represent restriction enzyme cutting site. SDM, site-directed mutagenesis.

**Table S4 References**


1. Wendoloski D, Ferrer C, Dyall-Smith ML. 2001. A new simvastatin (mevinolin)-resistance marker from *Haloarcula hispanica* and a new *Haloferax volcanii* strain cured of plasmid pHV2. Microbiology 147:959-64.

2. Allers T, Ngo HP, Mevarech M, Lloyd RG. 2004. Development of additional selectable markers for the halophilic archaeon *Haloferax volcanii* based on the *leuB* and *trpA* genes. Appl Environ Microbiol 70:943-53.

3. Humbard MA, Zhou G, Maupin-Furlow JA. 2009. The N-terminal penultimate residue of 20S proteasome α1 influences its $N^{\alpha}$ acetylation and protein levels as well as growth rate and stress responses of *Haloferax volcanii*. J Bacteriol 191:3794-803.

4. Reuter C, Uthandi S, Puentes J, Maupin-Furlow J. 2010. Hydrophobic carboxy-terminal residues dramatically reduce protein levels in the haloarchaeon *Haloferax volcanii*. Microbiology-SGM:248-255.

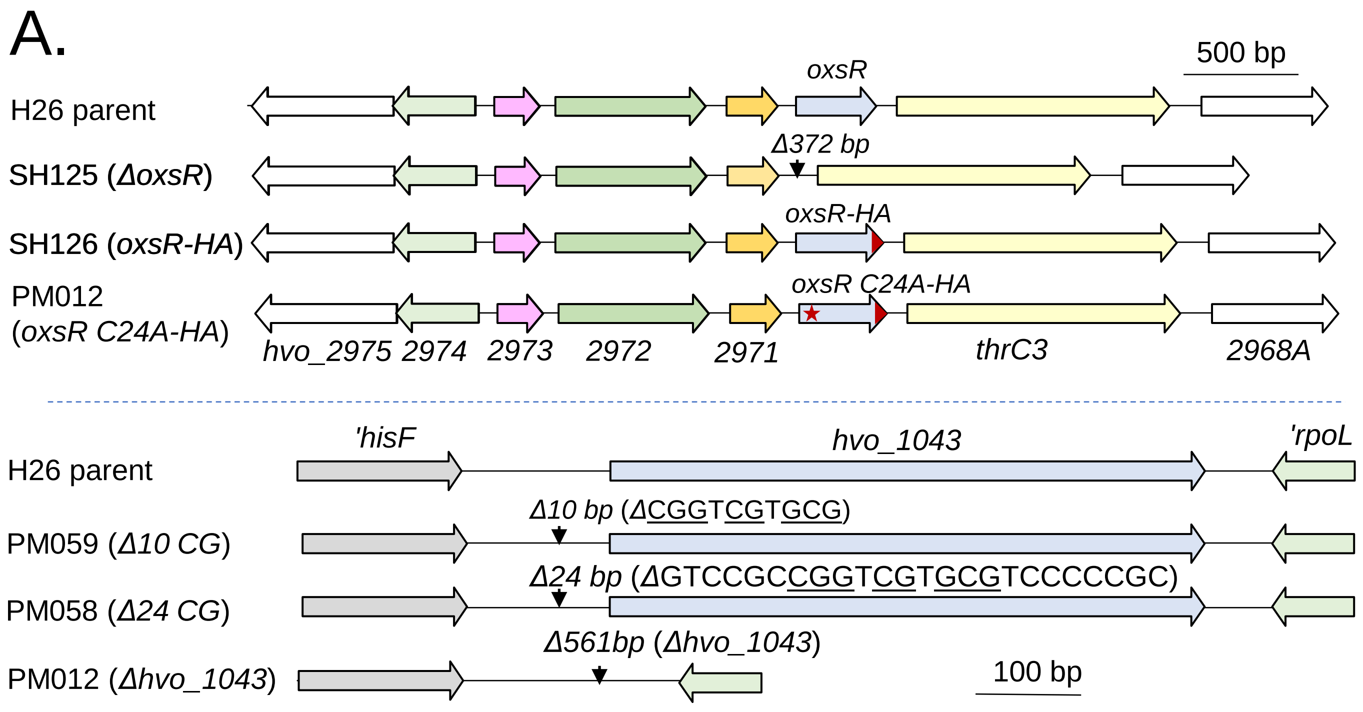


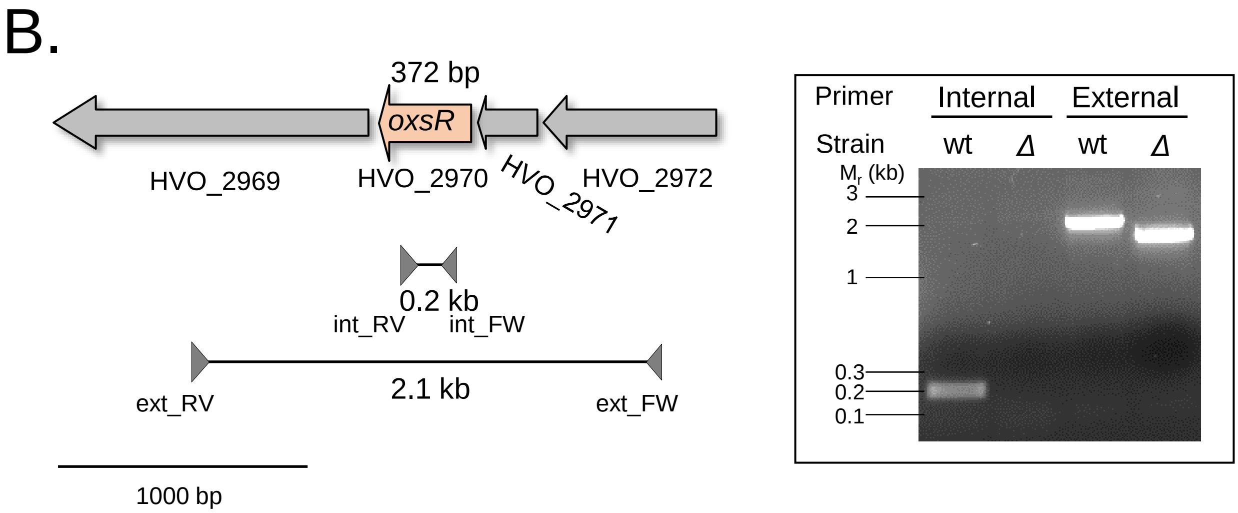


**Figure S1.** *H. volcanii* mutant strains generated in this study including SH125, SH126, PM012, PM057 and PM058, PM059. A. Genomic neighborhood of the mutant strains. Arrows represent the open reading frames with associated gene locus tag numbers indicated below. Triangle represents the site of the genomic deletion. Bar, scale 500 and 100 bp as indicated. B. Strategy used for markerless deletion as exemplified by generation of SH125. PCR was applied to confirm the *oxsR* gene deletion. A target gene specific primer set (lane 1 and 2) and an external primer set (680 – 970 bp flanking region from the target gene, lane 3 and 4) were used to confirm the *oxsR* gene deletion. Lane 1 and 3, gDNA from wt *Hfx. volcanii*; lane 2 and 4, gDNA from SH125.

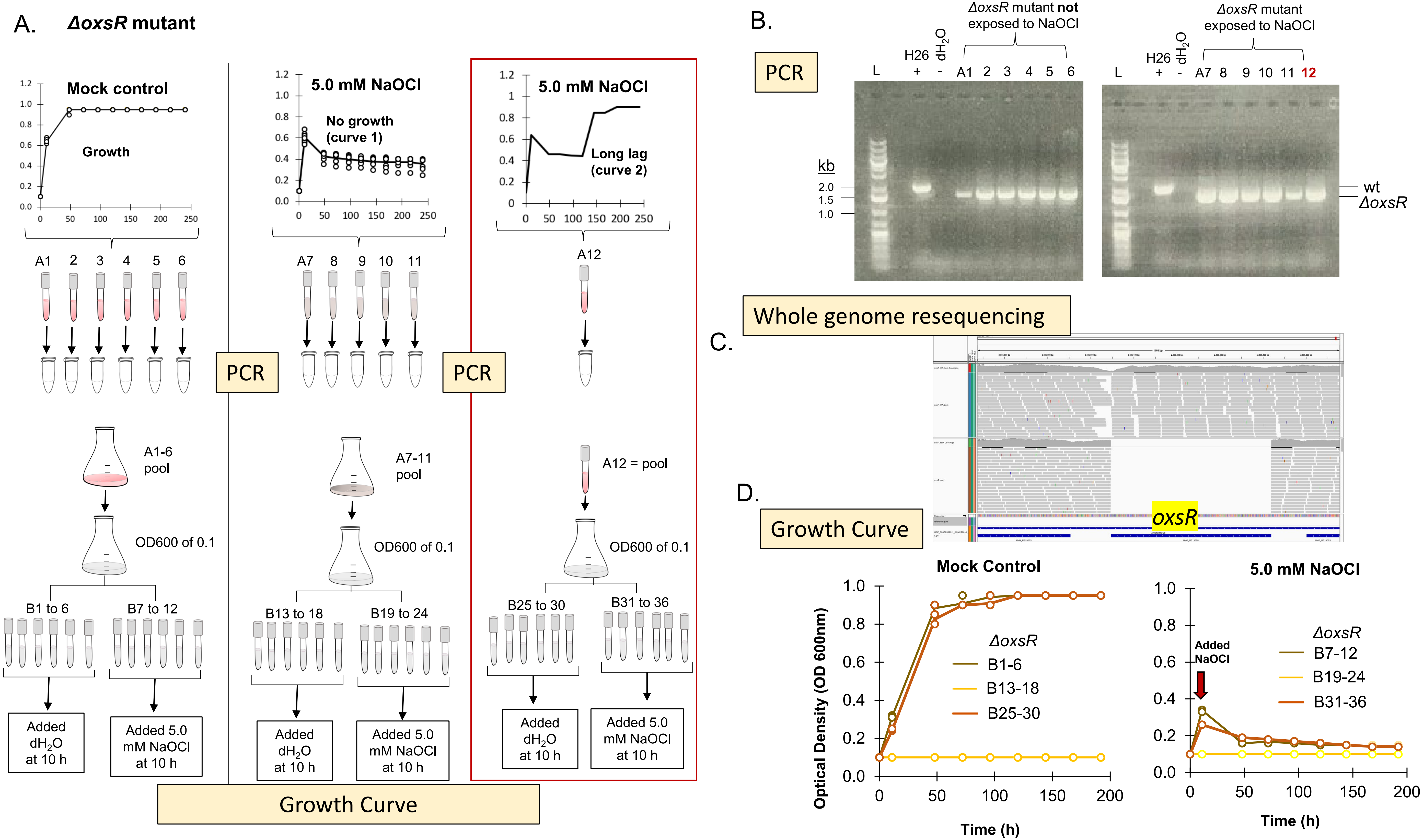


**Figure S2**. Further analysis of the *ΔoxsR* mutant. A) Experimental strategy used for analysis of the *ΔoxsR* mutant. All 12 tubes of experiment 2 were examined by PCR for retention of the *ΔoxsR* mutation. In addition, tubes A1-6, A7-11, and A12 were treated as three independent groups for analysis of hypochlorite sensitivity. Tubes A1-6 were pooled, tubes A7-11 were pooled, and tube A12 was retained as an independent sample The samples were diluted to an OD600 of 0.1 in fresh GMM and incubated at 42 °C with angled rotation for aeration as described in methods. After 10 h, the samples were treated with a mock control or NaOCl (5 mM) as indicated. Growth was monitored at OD600. B) PCR analysis to assess stability of the *ΔoxsR* mutant. Lanes: L, DNA standard (GeneRuler 1 kb Plus DNA Ladder, Thermofisher); *ΔoxsR* mutant (A1-12); parent (H26, +); nuclease free water (d$H_2O$, -). For PCR, cell culture (5 µl) was mixed with 30 µl of nuclease-free water and boiled for 10 min. Samples were centrifuged for 5 min at 13,000 × *g* and stored at -20°C for 4 days. Tubes were thawed on ice and centrifuged (5 min at 13,000 × *g*) and transferred (1 µl) for use as template in a 15 µl PCR reaction with primer pair 5/6 (outside the deletion plasmid). C. Whole genome resequencing indicates that OxsR was deleted from all copies of the genome (no reads are detected). Integrated Genomics Viewer (IGV) image shows sequencing reads (grey) of H26 parent strain (above) vs. Δ*oxsR* deletion strain (below). Blue lines in bottom track show the location of the genes in the locus. D. Growth curve of the *ΔoxsR* mutant cultures previously exposed to 5.0 mM NaOCl. See experimental strategy in panel A for tube numbering B1-36.

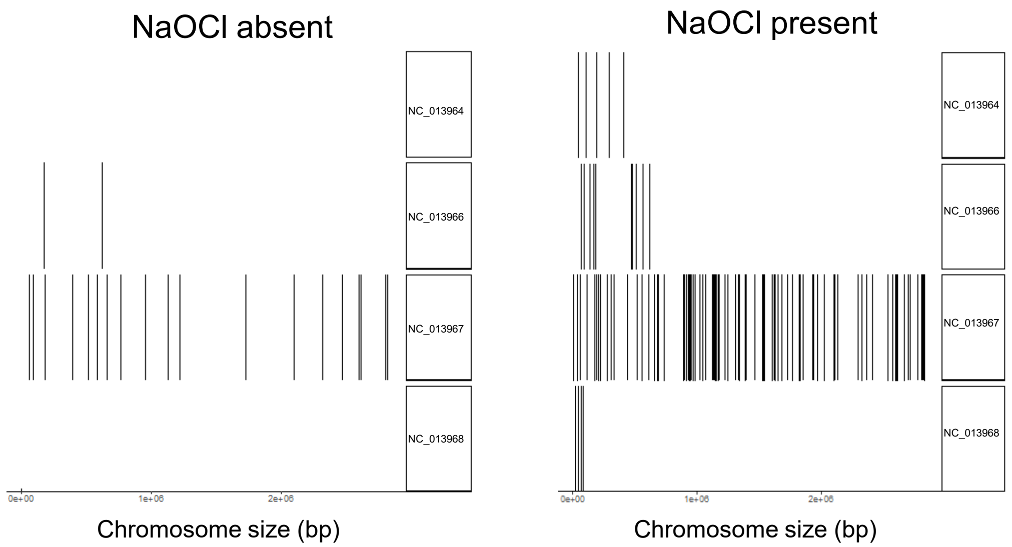


**Figure S3.** Peak loci in chromosome and plasmids by ChIP-seq analysis in the absence (left) or presence (right) of oxidative stress. NC_013967, main chromosome (2.85 Mb); NC_013968, pHV1 (0.09 Mb); NC_013965, pHV2 (0.01 Mb); NC_013964, pHV3 (0.44 Mb); NC_013966, pHV4 (0.64 Mb).

A.

**Network Stats**

| | | | |
|---|---|---|---|
| number of nodes: | 48 | expected number of edges: | 13 |
| number of edges: | 30 | PPI enrichment p-value: | 4.8e-05 |
| average node degree: | 1.25 | | |
| avg. local clustering coefficient: | 0.275 | | |

*your network has significantly more interactions than expected (what does that mean?)*

**Functional enrichments in your network**

*explain columns*

| GO-term | description | count in network | strength | false discovery rate |
|---|---|---|---|---|
| **Molecular Function (Gene Ontology)** | | | | |
| GO:0015036 | Disulfide oxidoreductase activity | 5 of 12 | 1.53 | 0.00080 |
| GO:0015035 | Protein disulfide oxidoreductase activity | 4 of 11 | 1.47 | 0.0053 |

| domain | description | count in network | strength | false discovery rate |
|---|---|---|---|---|
| **Protein Domains and Features (InterPro)** | | | | |
| IPR017937 | Thioredoxin, conserved site | 3 of 4 | 1.78 | 0.0374 |
| IPR036249 | Thioredoxin-like superfamily | 5 of 25 | 1.21 | 0.0321 |

B.

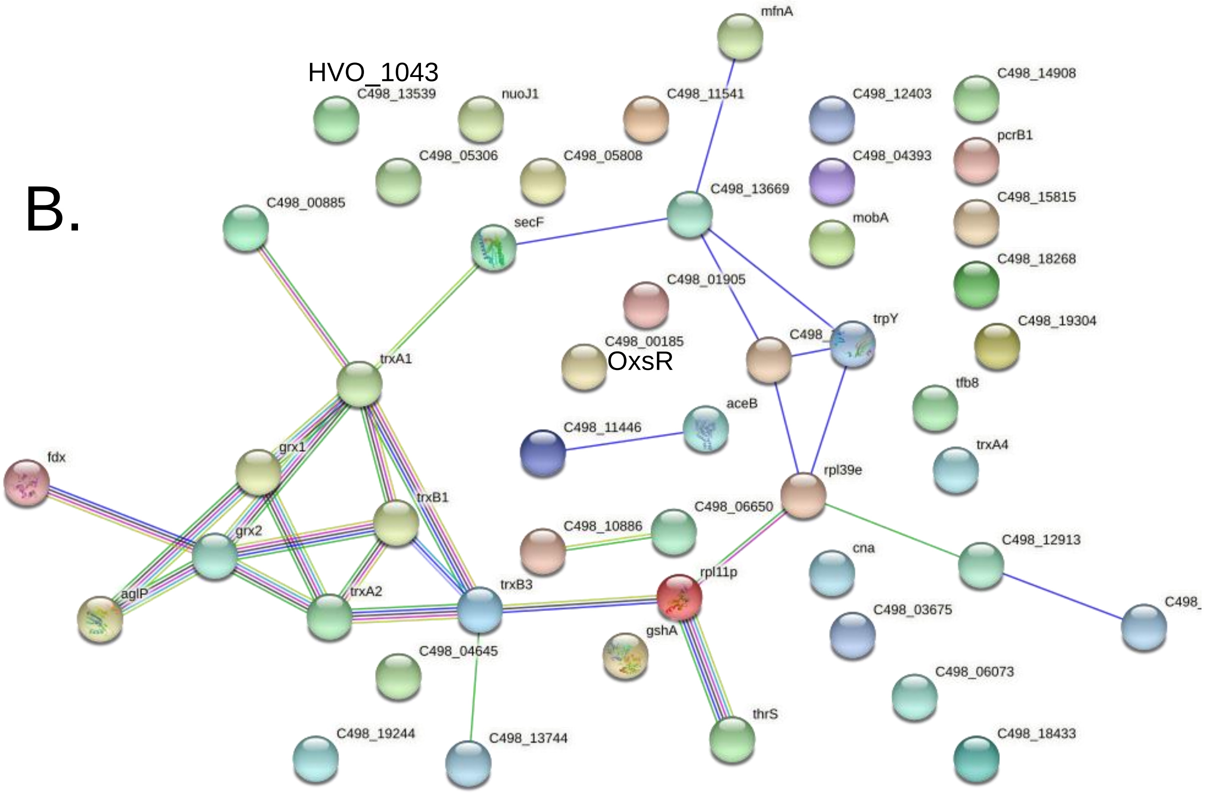


**Figure S4.** Functional enrichment of gene homologs associated with sulfur relay systems by STRING 11.5 analysis. OxsR and proteins encoded by genes immediately adjacent to OxsR-bound regions detected by ChIP-seq analysis analyzed by STRING (11.5) in tabulated (A) and node/edge configuration (B). Edges colored as: green line, neighborhood evidence; blue line, cooccurrence evidence; purple line, experimental evidence; yellow line, textmining evidence; light blue line, database evidence; black line, coexpression evidence. See **Table S3** for data input.

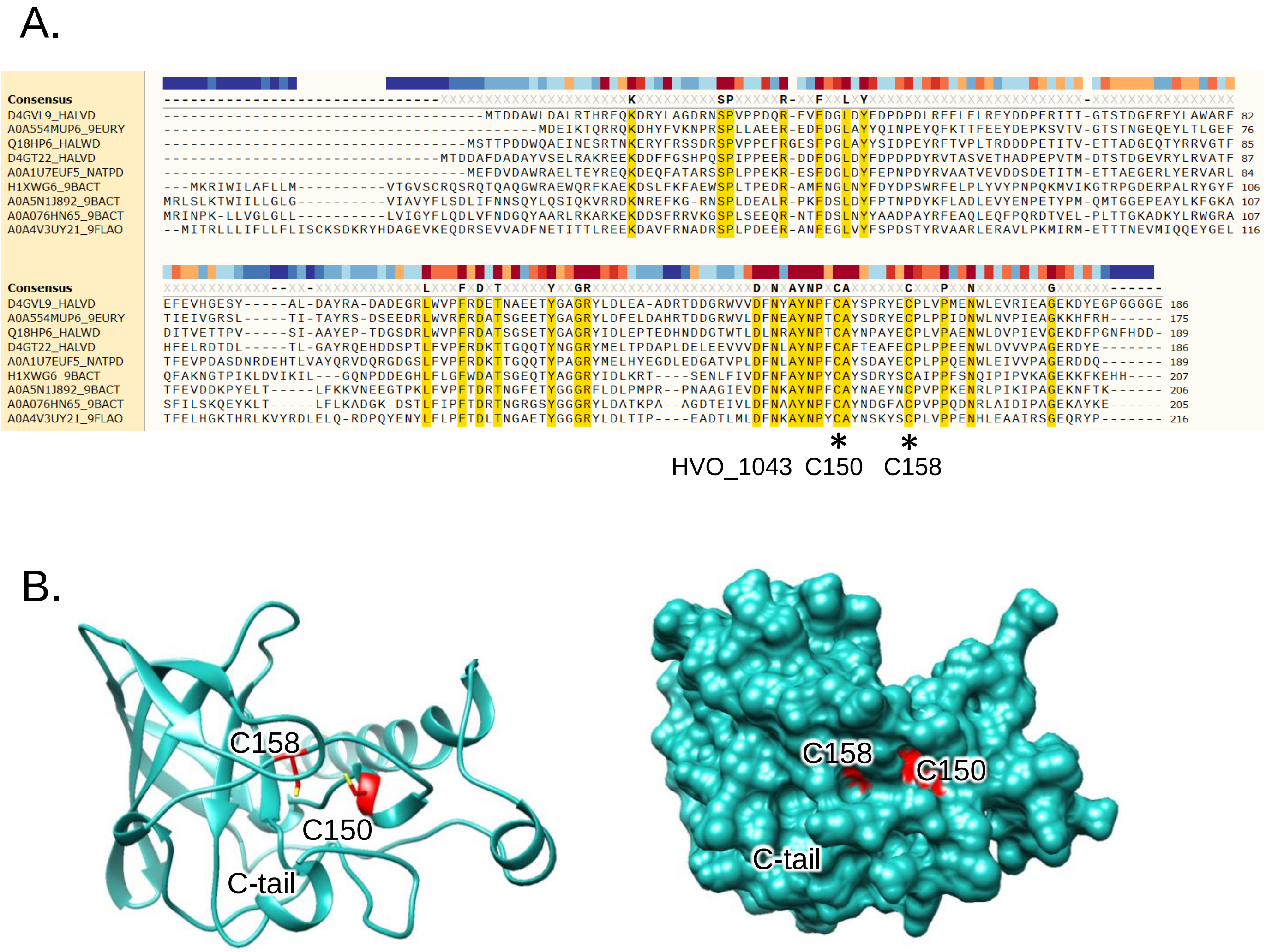


**Figure S5.** DUF1684 family protein HVO_1043 and its conserved $CX_7C$ motif suggest a role in thiol chemistry. A) Multiple amino acid sequence alignment of representative DUF1684 family members. UniProt reference numbers to left (D4GVL9 corresponds to HVO_1043). Orange highlight, residues > 95% sequence threshold. Bar above alignment colored red to blue, indicates regions of high to low sequence conservation, respectively. B) 3D-structural model of HVO_1043 represented as a ribbon (left) and surface (right) diagram. The C-terminal tail (C-tail) and conserved cysteine residues (C150 and C158) are indicated. HVO_1043 (96% of the sequence, 178 residues) was modeled with 100% confidence by the single highest scoring template: the NMR solution structure of *Haloarcula marismortui* rrnAC0354 (PDB: 2LNU). The NMR solution structure of *Halobacterium salinarium* VNG_0733h (PDB: 2LOK*)* was an additional high scoring (100% confidence) template.

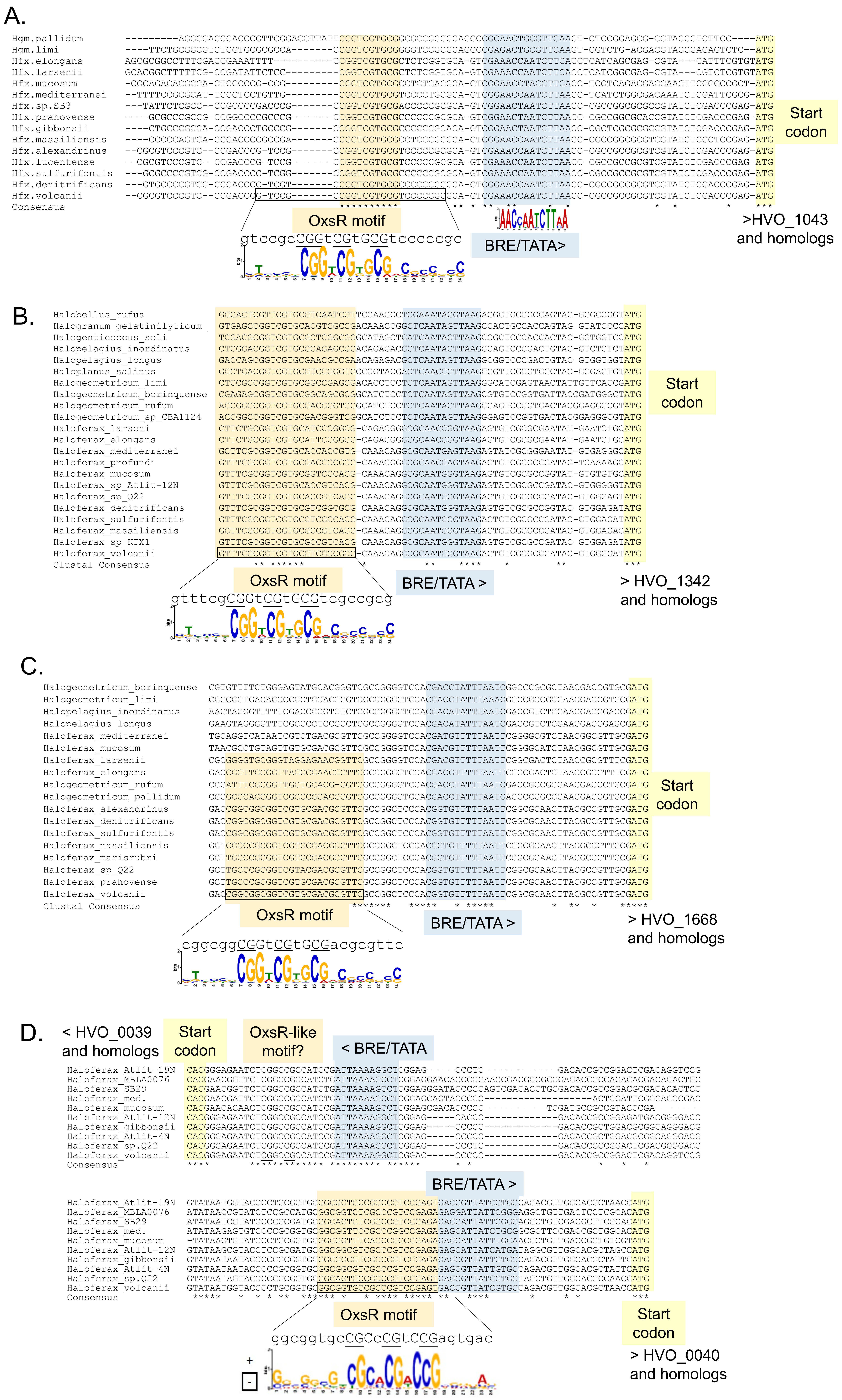


**Figure S6**. DNA motif common to OxsR bound regions and its location in relationship to the promoter consensus sequence element in the 5' region of select gene homologs identified by ChIP-seq analysis. A) HVO_1043, B) HVO_1342, C) HVO_1668, and D) HVO_0039 and HVO_0040 and their homologs were included in this analysis.

A

28% amino acid identity

```
HVO_1360 --MASSMVEYL-QSDMECEGLLECLHGLKQLDRRCFEVLVETDDRLTVDEVAEAVERERS 57
OxsR     MADAPDMGELMETEDPNFGQVLACVFGIQSHESRTYLALLDN-PGSTVAELAEVLDRDRS 59
            * .* * :  .* :   :* *:.*::. : * : .*::.    ** *:**.::*:**
HVO_1360 TAYRSIQRLLQAGLIQKQQVNYEHGGYYHVYHPTDPNEVADDMQRMLNDWYAQMGTLIQE 117
OxsR     NVNRSLTTLLDKGLTERKRRLLDPGGYVYQYTATPLPEAKEMMHAALDEWAEDVHARIDA 119
         .. **:  **: ** ::::   : *** : *  *   *. : *:  *::*  :: : *:
HVO_1360 FRDKYDEKIVPAE 130
OxsR     FGES--------- 123
         * :.
```

B

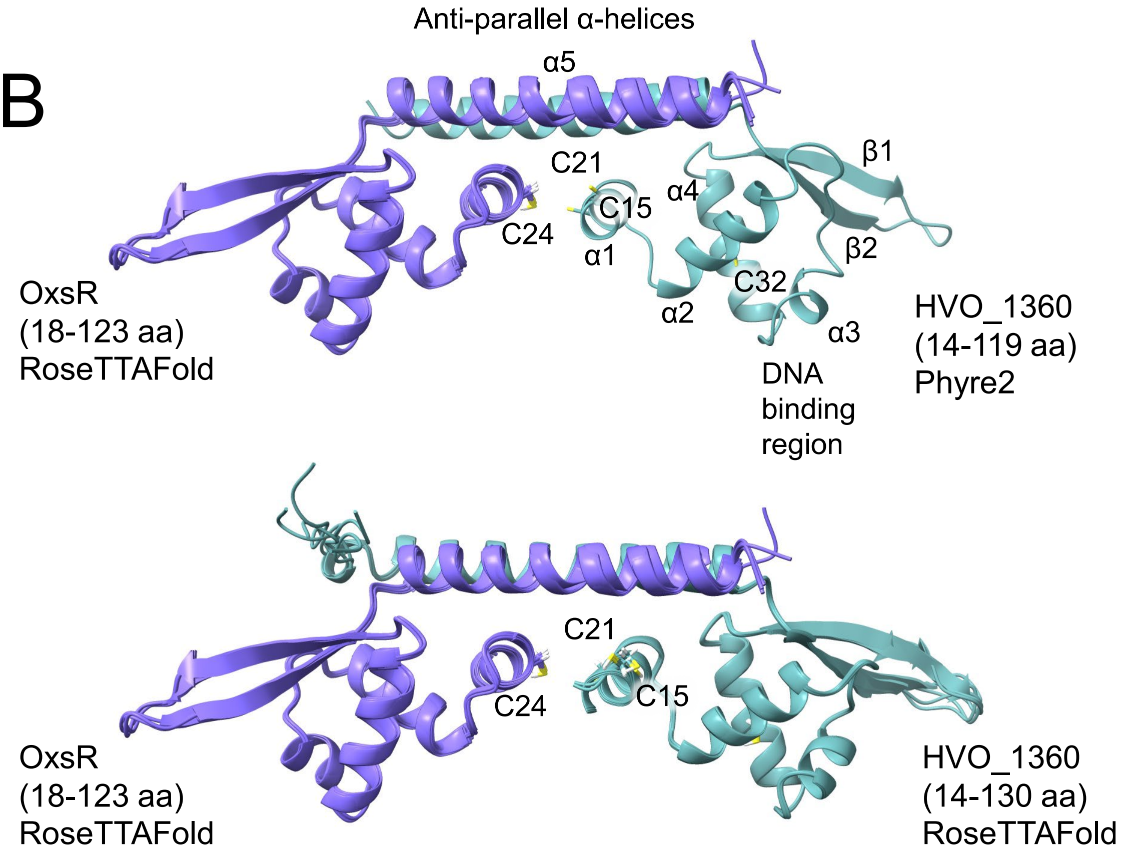


**Figure S7.** TrmB family proteins HVO_1360 and OxsR (HVO_2970) are structurally related. A) Amino acid sequence alignment of HVO_1360 and OxsR with identical (*), functionally similar (. or :), cysteine (red), and predicted DNA binding residues (purple) indicated. B) 3D-structural models of HVO_1360 (cadet green) and OxsR (purple). The monomeric 3D-structures were oriented as a heterodimer using *Sulfolobus acdiocaldarius* AbfR2 (PDB: 6CMV) as a scaffold. Models are presented as ribbon diagrams with cysteine residues as stick diagram and were performed by RoseTTAFold and Phyre2 as indicated. The N-terminal residues of OxsR (1-17 aa) and HVO_1360 (1-13 aa) which appeared variable in the 3D-models were manually removed. The OxsR and HVO_1360 Phyre2 models were related at an RMSD of 2.212 Å across all 106 atom pairs.
